# Context-dependent limb movement encoding in neuronal populations of motor cortex

**DOI:** 10.1101/588129

**Authors:** Wolfgang Omlor, Pia Sipilä, Anna-Sophia Wahl, Henry Lütcke, Balazs Laurenczy, I-Wen Chen, Lazar T. Sumanovski, Marcel van ’t Hoff, Philipp Bethge, Fabian F. Voigt, Martin E. Schwab, Fritjof Helmchen

**Affiliations:** Laboratory of Neural Circuit Dynamics, Brain Research Institute, University of Zurich, Zurich, Switzerland.; Laboratory of Neural Regeneration and Repair, Brain Research Institute, University of Zurich, and Dept. of Health Sciences and Technology, ETH Zurich, Zurich, Switzerland.; Neuroscience Center Zurich, University of Zurich and ETH Zurich, Zurich, Switzerland.

## Abstract

Neuronal networks of the mammalian motor cortex (M1) are important for dexterous control of limb joints. Yet it remains unclear how encoding of joint movement in M1 networks depends on varying environmental contexts. Using calcium imaging we measured neuronal activity in layer 2/3 of the mouse M1 forelimb region while mice grasped either regularly or irregularly spaced ladder rungs during locomotion. We found that population coding of forelimb joint movements is sparse and varies according to the flexibility demanded from them in the regular and irregular context, even for equivalent grasping actions across conditions. This context-dependence of M1 network encoding emerged during learning of the locomotion task, fostered more precise grasping actions, but broke apart upon silencing of projections from secondary motor cortex (M2). These findings suggest that M2 reconfigures M1 neuronal circuits to adapt joint processing to the flexibility demands in specific familiar contexts, thereby increasing the accuracy of motor output.

## INTRODUCTION

In everyday life we have to generate dexterous movements of limb joints to purposefully interact with variable environments. The mammalian primary motor cortex (M1) is known to contribute to control of dexterous limb movements^1–8^ and gait modifications^9–11^. Moreover, M1 has recently been shown to have a pivotal function in learning non-dexterous movement sequences in rats^12^, and has been suggested to be necessary for progressing though the steps of learned skilled forelimb movements in mice^7^. Still, the principles of its operation – such as the representation of movements in the M1 microcircuit – remain poorly understood^13–19^. Various studies linked changes of movement parameters to changes of neuronal M1 activity during ongoing motor actions. For example, neuronal activity in M1 was found to control various movement variables including direction, force, speed, end-posture and individual joint angles as well as muscle activity^14,15,17,18,20–27^. However, in addition to representing features of the ongoing movement itself, neuronal activity in M1 may also represent general demands of the environmental setting, within which the movement is executed. These demands could be regarded as “meta-variables” that adjust activity of neuronal circuits in M1 in addition to the movement variables that characterize the ongoing motor action itself. The representation of a specific limb movement may be flexibly modulated according to certain principles when the same motor action is executed in different environmental contexts. Such context-dependent modulation of M1 neuronal encoding of limb movements has been scarcely investigated so far, but the existence for context impact on motor control is assumed for example by recent studies of the mirror neuron network^28–30^.

Accurate motor control in a given environmental context requires integration of contextual information with processing of specific sensory stimuli^31^. Layer 2/3 (L2/3) neurons in M1 receive inputs from sensory areas^32,33^ as well as from the secondary motor cortex (M2)^34–36^, which is thought to convey context information for motor processing^37–39^. Given this pattern of afferent inputs and their excitatory output to L5 neurons in M1^40,41^, L2/3 neurons are well positioned to optimize motor commands by integrating contextual and sensory information, consistent with a pivotal role in the refinement of motor actions^42–44^.

Here we hypothesized that the degree of flexibility that is demanded from an individual limb joint in a specific environment is a key contextual parameter that affects joint movement representation in L2/3 of M1. We applied two-photon calcium imaging to record L2/3 neuronal population activity in M1 while mice performed distinct forelimb grasping actions in order to move on either regularly or irregularly spaced ladder rungs. Compared to the regular pattern, the irregular pattern represents a different environmental context that demands more flexible use of several limb joints, for example of the proximal shoulder joint. Lesion studies demonstrated that the motor cortex is required for accurate forelimb movements on the regular and in particular on the irregular pattern in mice as well as rats^3,45,46^. We show that joint movements are differentially encoded in L2/3 neuronal networks of M1 in the regular and irregular context, respectively, even for motor actions with matching kinematic profile, and that encoding strength increases if a higher flexibility is demanded and vice versa. Furthermore, we demonstrate that this ‘context-dependent coding of flexibility demands’ emerges when the animal is familiarized with the distinct contexts and learns to interact with the respective situations. Finally, using chemogenetic silencing we show that context-dependent modulation of L2/3 neuronal representation in M1 entails more precise limb movements and requires input from M2.

## RESULTS

### Grasping behavior on regular and irregular ladder rung wheels

To enable calcium imaging of cortical neurons in awake, head-fixed mice under different contextual conditions, we customized the rung ladder test for rodents^3,45,46^. We built two ladder wheels (23-cm diameter), one with rungs at constant 1-cm spacing (‘regular’ wheel), the other one with rungs placed at distances varying unpredictably between 0.5 to 3 cm (‘irregular’ wheel; 0.5-cm steps; 1.68 ± 0.56 cm spacing, mean ± s.d.; Fig. 1a; Online Methods). We trained mice to perform skilled locomotion on these ladder wheels, with all mice reaching saturating forelimb performance scores^3,45^ for both conditions within 8 training days (Supplementary Fig. 1; Online Methods). Hence, days 9-12 were defined as ‘expert phase’. Using high-speed videography, we tracked kinematic changes of the shoulder, elbow, wrist, and finger base joints for the right forelimb of trained mice for each ‘run’, during which mice covered a predefined distance with a continuous sequence of forelimb grasps (Fig. 1a,b; 11.1 ± 3.5 grasps per run [24.7 ± 6.4 runs] and 8.8 ± 2.4 grasps per run [25.7 ± 6.1 runs] for regular and irregular wheel, respectively; mean ± s.d.; n = 7 mice). The distributions of reaching distance (RD) and grasp duration (GD) were similar and not significantly different for both types of wheels across mice (Fig. 1c; ROC-AUC = 0.55 and 0.64 for RD and GD, respectively, comparing regular and irregular distributions using the ‘area under the curve’ (AUC) of the receiver operating characteristic (ROC); see Online Methods; pooled across all 7 animals during the expert phase).

**Figure 1.**
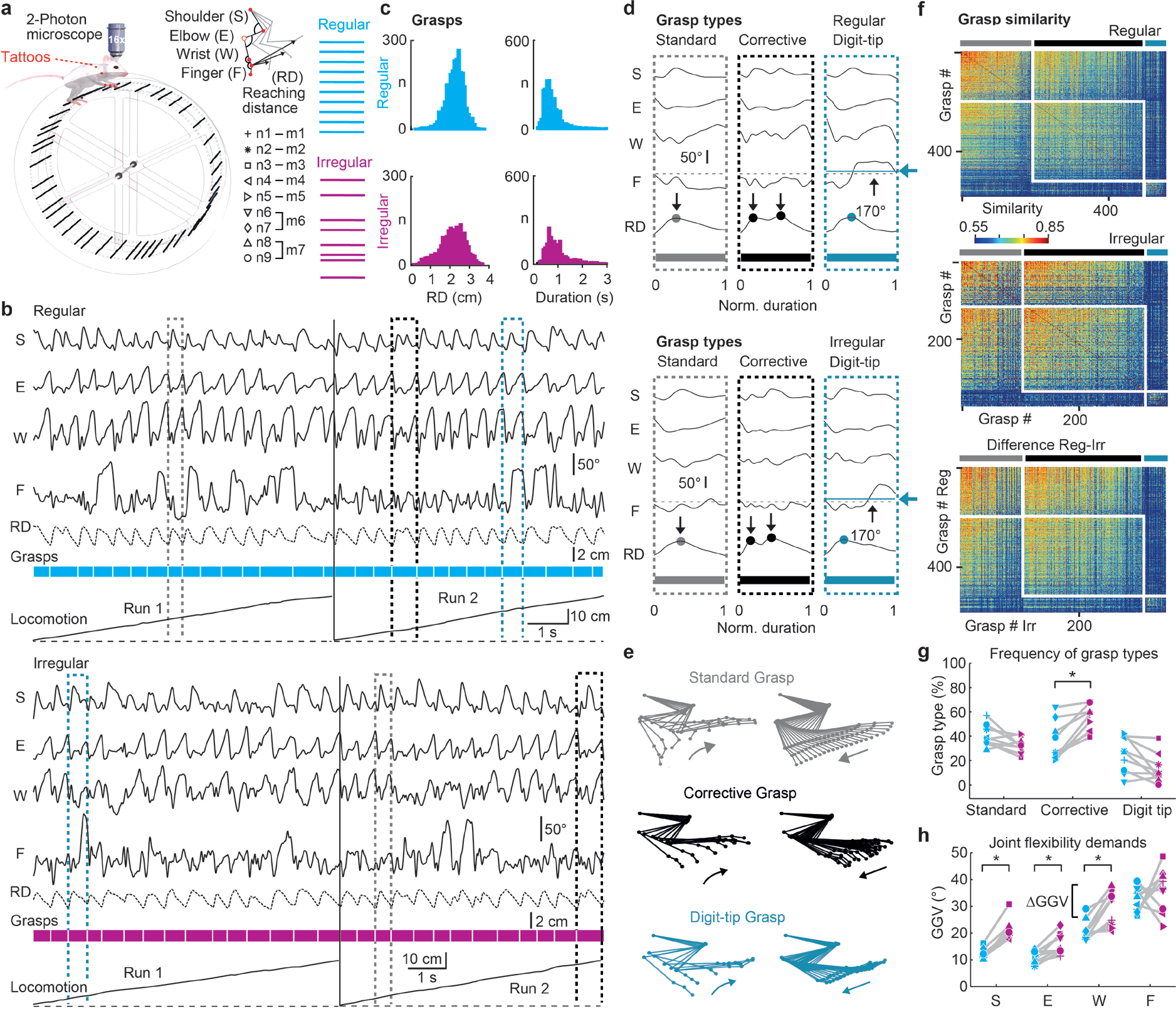
Quantitative analysis of forelimb grasps during skilled locomotion on regular and irregular ladder wheels. (**a**) Schematic of setup with head-fixed mouse on top of a ladder wheel and below a two-photon microscope, moving across rungs with regular (cyan) or irregular (magenta) spacing (here the irregular wheel is shown). 9 neuronal networks in M1 L2/3 were recorded (n6 and n7 as well as n8 and n9 belong to mice 6 and 7, respectively). A video camera tracks tattoos on the right forelimb (red dots: scapula, shoulder, wrist, finger base joint, and digit tip; the elbow position is calculated assuming fixed shoulder-to-elbow and elbow-to-wrist distances). Joint angle changes in shoulder (S), elbow (E), wrist (W) and finger base (F), as well as reaching distance (RD) are quantified. (**b**) Time course of forelimb joint angles and RD during two concatenated example runs on the regular (top) and irregular (bottom) wheel with individual grasps indicated. Three prototypical grasps are highlighted: Standard (grey), corrective (black) and digit-tip grasp (dark turquois). (**c**) Histograms of maximal RD during each grasp and grasp duration for both conditions, pooled across all 7 mice. Inlays display the value of the area under the ROC-curve for the regular and irregular distribution. (**d**) Kinematic profile of joint angles and RD for examples of standard (grey), corrective (black), and digit-tip (dark turquois) grasps during both conditions (grasps marked in (b) with duration normalized). Dots on RD traces indicate the number of reaching cycles during the grasp; black horizontal dashed lines mark 170° threshold for mean finger extension, which is exceeded only in digit-tip grasps as indicated by the dark turquois dotted line and arrow). Time scale is normalized from start (0) to end (1) of grasps. (**e**) Representative stick-figure plots of limb kinematics for the three principal grasp types (left: reaching phase, right: pulling phase, same color code as in (b) and (d)). (**f**) Grasp-similarity matrices of one example mouse for the regular and irregular condition as well as for the difference between both conditions, sorted according to the classification in standard (grey), corrective (black) and digit-tip (dark turquois) grasps. Matrices are sub-sorted according to similarity values. (**g**) Fraction of grasps types on the regular and irregular wheel for all 9 neuronal networks. (**h**) Grasp-to-grasp variability of each joint on the regular and irregular pattern for all 9 neuronal networks. Asterisks indicate *P* < 0.05 (paired t-test, P-value adjusted according to HB).

Across mice and conditions we identified three salient grasp types based on the temporal profile of the reaching movement and the mean finger extension during each grasp (Fig. 1d,e; see Online Methods for details of classification criteria; **Supplementary Video 1**): ‘*Standard*’ grasps consisted of a single motion cycle including reaching phase, correct placement of the forepaw on the rung, closure of the paw, and a terminal pulling phase; ‘*corrective*’ grasps were characterized by one or multiple corrective movements after the initial reaching action until the forepaw optimally hit the targeted rung with its palm and the subsequent pull occurred; and finally, ‘*digit-tip*’ grasps, during which the targeted rung was hit with the digit tips rather than with the palm of the hand, causing pronounced extension of the finger base joint and necessitating dexterous finger control to avoid a slip and to finish the pull. This classification was also reflected in similarity matrices for grasp pairs, which we calculated using the mean Euclidian distance of grasp trajectories in 4-dimensional joint angle space (trajectories were time-warped to account for different grasp durations; Online Methods). Digit-tip grasps formed a cluster clearly separate from the other grasp types under both conditions whereas the distinction between standard and corrective grasps, which showed considerable diversity, was less obvious (Fig. 1f). The fraction of corrective grasps was significantly higher for the irregular compared to the regular wheel (Fig. 1g; *P* = 0.0063, *t* = −5.1651; paired t-test with *P* adjusted according to Holm-Bonferroni, HB; df = 6; n = 7; Online Methods). To quantify the flexibility demands for each forelimb joint in the regular and irregular context, respectively, we also quantified the grasp-to-grasp variability (GGV) of joint motion by calculating the mean amplitude variation from one grasp to the next (Online Methods). For shoulder, elbow and wrist, GGV was significantly higher for the irregular compared to the regular rung pattern (Fig. 1h; *P* = 0.0004, t = −9.2052, *P* = 0.0121, t = −4.5128, and *P* = 0.0225, t = −3.6083, respectively; paired t-test with HB adjustment; df = 6; n = 7). In contrast, GGV for the finger joint was not significantly different between conditions (Fig. 1h; *P* = 0.4769, t = −0.7585). These results indicate that shoulder, elbow and wrist movements consistently require more flexible recalibration from grasp to grasp on the irregular wheel compared to the regular wheel. In contrast, the flexibility demands of finger-base movements do not show a consistent dependency on regular versus irregular context.

### Calcium imaging in identified M1 forelimb area during grasping

To consistently localize the forelimb region in M1 for subsequent calcium imaging we used transgenic mice that express channelrhodopsin-2 (ChR2) under the thy1 promoter in cortical L5 neurons^47,48^ and in addition virally expressed in M1 the genetically encoded calcium indicator yellow-cameleon Nano140 (YC-Nano140)^49–51^. To identify the M1 forelimb area, we performed optogenetic motor mapping^47^ by scanning a blue laser spot across M1 to stimulate localized networks of L5 neurons and concurrently measuring light-evoked joint movements of the contralateral forelimb using video monitoring (Fig. 2a,b; Online Methods). Light stimulation caused forelimb muscle contractions and induced flexion or extension of individual joints within confined but overlapping regions (Fig. 2b). Within this forelimb region, all mice featured a central spot, in which the combined evoked movement of all joints was maximal (Fig. 2b,c; Online Methods). In 6 of the 7 animals, activation of this ‘forelimb focus’ elicited retroversion of the shoulder joint, flexion of elbow and wrist joints as well as extension of finger base joints, resembling the initial phase of a grasp (Fig. 2c).

**Figure 2.**
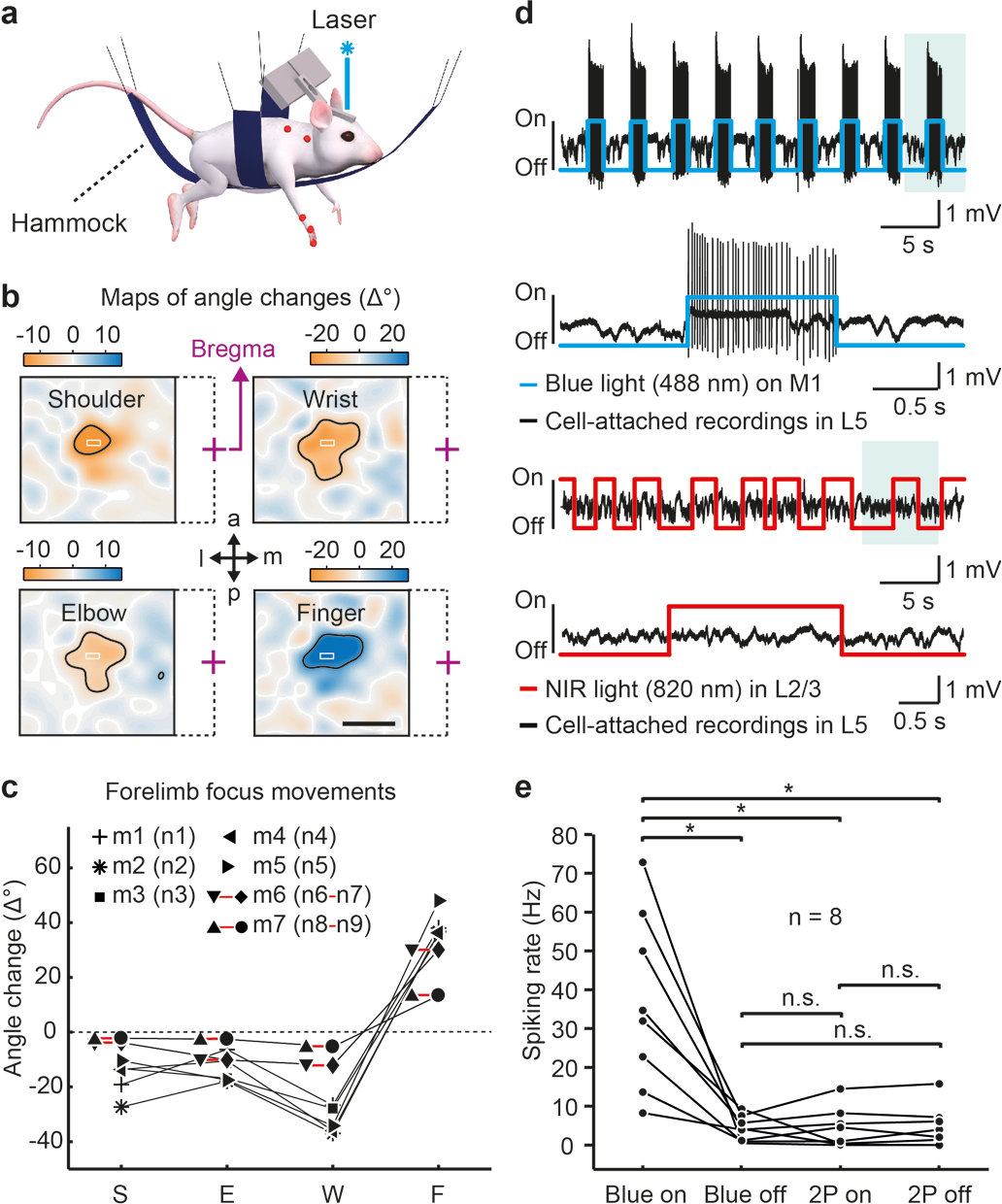
Optogenetic M1 mapping and electrophysiological recording of L5 neurons during two-photon laser scanning in transgenic ChR2 mice. (**a**) Schematic of motor mapping setup with an anesthetized mouse hanging in a hammock, limbs dangling free. Blue laser light randomly scanned across M1 evoked forelimb joint angle movements that were monitored with a camera. (**b**) Maps of evoked joint angle changes for an example mouse. Negative values correspond to flexion (orange, corresponding to retroversion in the shoulder joint), positive values to extension (blue, corresponding to anteversion in the shoulder joint). Laser-stimulation elicited responses in shoulder (area 1.12±0.19 mm^2^), elbow (1.29±0.27 mm^2^), wrist (1.16±0.32 mm^2^) and finger base joints (1.03±0.13 mm^2^; thresholded at 50% of maximal response in each joint, black contour). The superimposed white rectangles indicate the selected “forelimb focus” area for subsequent calcium imaging (purple cross = bregma, dashed lines indicate the affiliation to the respective map; a = anterior; p = posterior; m = medial; l = lateral). Scale bar 1 mm. (**c**) Movement amplitude of all joint angle changes in the seven mice induced by optogenetic stimulation in the forelimb focus. (**d**) Upper part: Cell-attached recording of a ChR2-expressing L5 neuron in M1 during repetitive application of blue 488-nm light. The evoked spiking pattern during one stimulation period (shaded area) is shown on expanded time scale below. Lower part: Cell-attached recording of the same L5 neuron during two-photon excitation laser scanning with near-infrared (NIR) light in L2/3, equivalent to the conditions used for L2/3 calcium imaging. The expanded view of one stimulation period below demonstrates the lack of two-photon excited spikes and extracellular voltage changes. (**e**) Pooled data for similar recordings in eight L5 neurons, indicated by black dots (neurons were not recorded from the same ChR2-mice that are shown in c). Whereas blue light stimulation induced strong spiking of L5 neurons, laser scanning in L2/3 with two-photon (2P) excitation light of 820-nm wavelength did not induce any detectable changes in the spiking rate of L5 neurons. Asterisks indicate significant differences with *P* < 0.05; paired t-test; Bonferroni-Holm corrected; n.s. (non-significant) means *P* > 0.05.

These spots were selected for two-photon calcium imaging of L2/3 neuronal populations. In one animal, the elbow joint showed extension rather than flexion in the stimulation spot of the strongest forelimb movement. In this animal, a different spot featuring the largest combination of shoulder retroversion, elbow and wrist flexion as well as extension of finger base joints like in the other animals was selected for two-photon calcium imaging (Fig. 2c). In control experiments using cell-attached recordings from ChR2-expressing L5 neurons, we verified that laser-scanning for two-photon imaging in L2/3 did not cause spurious spiking activity in L5 (Fig. 2d,e; Online Methods). We then measured fluorescence signals of YC-Nano140-expressing L2/3 neurons while expert mice engaged in skilled locomotion on the regular and irregular wheel, respectively (9 imaging areas with the same neuronal network recorded in both contexts; 493 cells in total; 7 mice; 54.8 ± 13.3 neurons per area; 195 to 220 µm below the pia; **Supplementary Video 2**). For running periods on both wheel types we extracted neuronal calcium transients from somatic regions of interests (ROIs). For further analysis we deconvolved calcium transients^49^ to infer the time course of instantaneous spiking rate changes (SR) (Fig. 3a-c; Online Methods). During runs, L2/3 activity was heterogeneous across the sampled neuronal subsets and temporally sparse, with occasional large calcium transients (>15% ΔR/R; >10 Hz SR) indicating bursts of action potentials. Some cells showed calcium transients that were strongly linked to salient finger movements as they occurred during digit-tip grasps in both conditions or to large shoulder movements on the irregular wheel (Fig. 3b,c; see Supplementary Fig. 2 for further cell examples).

**Figure 3.**
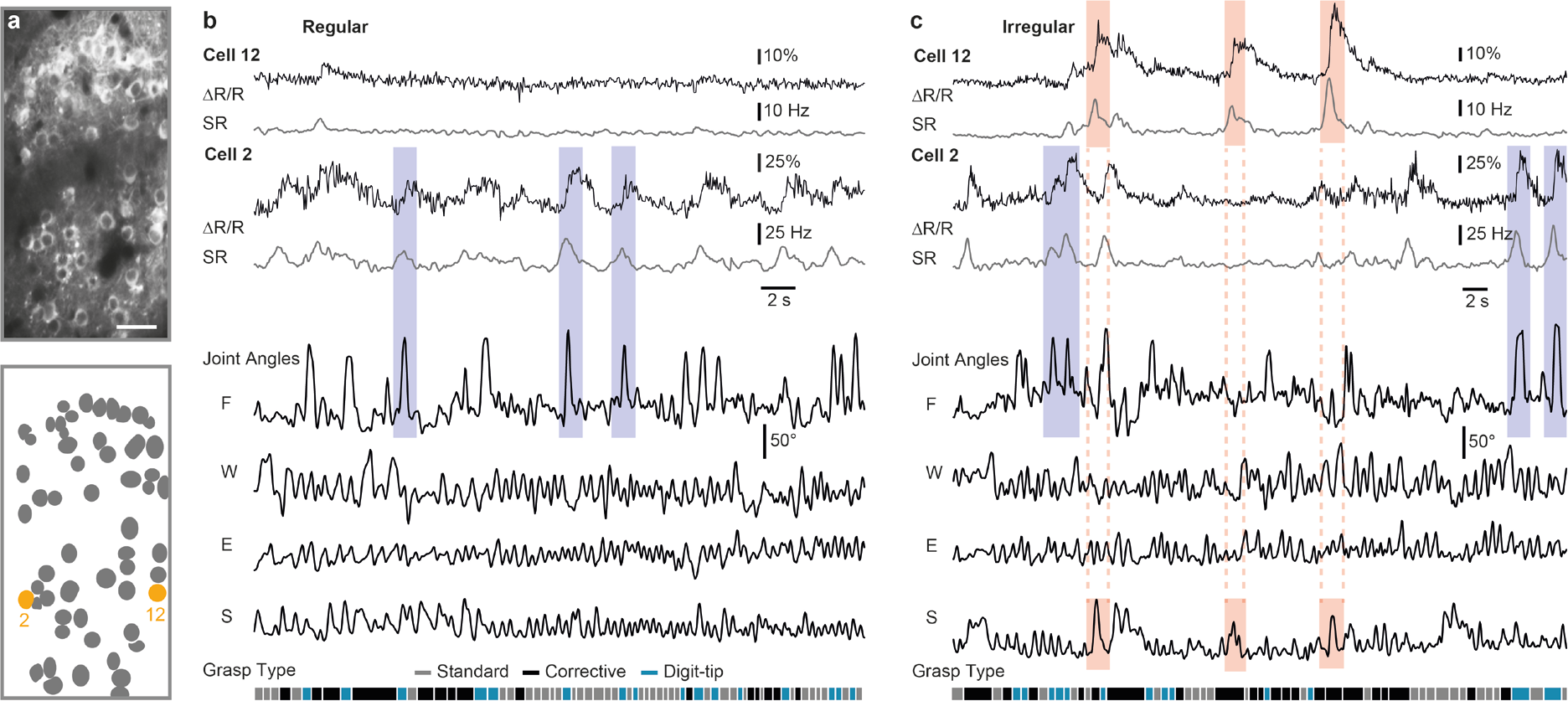
Neuronal activity in M1 L2/3 and simultaneous forelimb joint movements during skilled locomotion. (**a**) Example two-photon image of L2/3 neuronal population (upper panel) imaged during skilled locomotion with the genetically encoded calcium indicator *YC-Nano140* and schematic of ROIs (lower panel) with two neurons marked in orange; Scale bar 50 µm. (**b**) Raw *YC-Nano140* traces (ΔR/R, thin black line) and deconvolved instantaneous spiking rates (SR, thick grey line) for the two example cells marked in (a) for the regular condition along with simultaneously recorded joint angles and classified grasp types. Three salient finger movements (light blue shaded areas) that correlate with neuronal activity are highlighted. (**c**) Same conventions as in (b) for the irregular condition. Three salient finger movements (light blue shaded areas) and 3 strong shoulder movements (light red shaded areas) as well as the simultaneous neuronal responses are highlighted.

### Differential encoding of joint movements according to contextual flexibility demands

As a simple first analysis we evaluated how the activity of individual neurons related to specific grasp types by averaging grasp-related calcium transients separately for each grasp type (Fig. 4a). A small but significant fraction of neurons exhibited grasp-related activity for digit-tip grasps (6.9% of all cells on the regular and 6.1% of all cells on the irregular wheel; Fig. 4b; above chance level of 0.003%; Online Methods). 0.2% of neurons in the regular context and 0% of cells in the irregular context showed activity correlated with standard and corrective grasps. Thus, M1 L2/3 neurons show pronounced activity mainly during digit-tip grasps, whose salient feature is the extensive movement of finger base joints. We therefore next asked to what extent single neurons predict the time course of finger movements, along with the time course of shoulder, elbow and wrist movements. We trained a random forest algorithm to predict each of the four joint angles based on the deconvolved calcium signals of individual neurons (see Online Methods). On the regular wheel, no neuron predicted elbow movements and very few individual neurons predicted shoulder (0.6%) and wrist (0.4%) movement (ROC-AUC between true and shuffled prediction ≥ 0.95, see Online Methods). On the irregular rung pattern, however, 3% of neurons significantly predicted shoulder, 0.4% elbow and 3% wrist movements (Fig. 4c, two pies on the left). Under both conditions, about a tenth of neurons significantly predicted finger movements, with a slightly higher fraction for the regular pattern (9% compared to 7% on the irregular pattern). Thus, when mice have learned to step on the irregular ladder, the number of M1 L2/3 neurons encoding shoulder, elbow and wrist motion increases compared to the regular ladder, at the expense of neurons encoding finger motion. As expected from the grasp-type-related analysis, a large fraction of the neurons showing significant activity during digit-tip grasps also significantly predicted finger-base movements in the regular (76%) and irregular condition (47%). Of the 493 recorded neurons, 5.5% significantly predicted a particular joint movement only in the regular and 8.75% only in the irregular context, indicating context-specific reconfiguration of neuronal circuits in M1 L2/3. 2.9% of the 493 neurons predicted a particular joint movement in both conditions and only 1.4% switched the prediction of a particular joint movement for the regular and irregular context (Fig. 4c, pie on the right).

**Figure 4.**
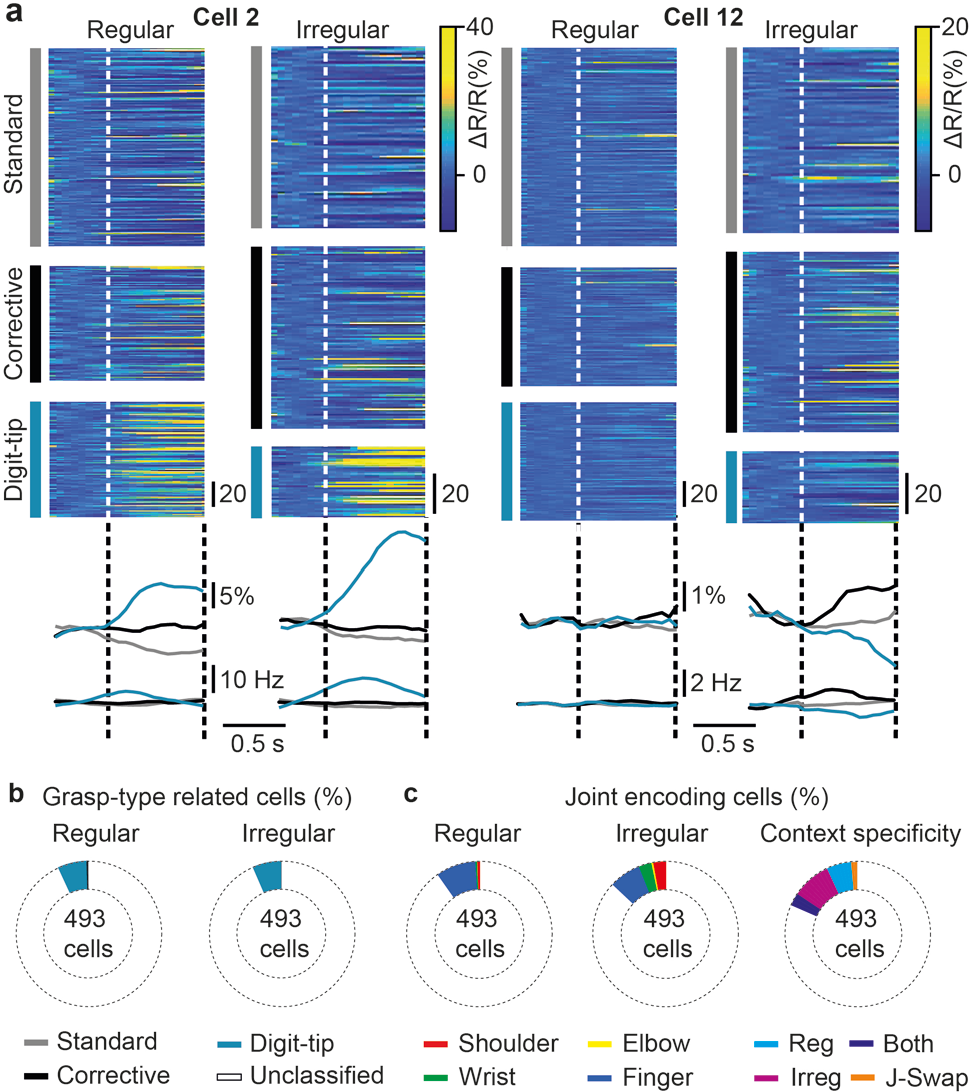
Responses of single neurons to grasp types and movements in individual joints. (**a**) Raw calcium (ΔR/R) traces for cell 2 (left) and cell 12 (right) from Fig. 3 during the three different grasp types and for both conditions. Corresponding average traces below for raw calcium traces (upper panel) and devonvolved spiking rate (SR) traces. (**b**) Percentages of the 493 recorded cells showing significant activity related to specific grasp types under both conditions (pooled across all 9 recorded neuronal networks). (**c**) Percentages of the 493 recorded neurons showing significant prediction of specific forelimb joint angle changes under both conditions (two pies on the left, pooled across all 9 recorded neuronal networks). In Figs. 3 and 4a, cell 2 is classified as finger-predictive and significantly active during digit-tip grasps; cell 12 is classified as shoulder-predictive. Pie on the right: Context specificity of significant joint-angle predictive cells from regular to irregular. Neurons significantly predicted a particular joint movement either only in the regular condition (‘Reg’, cyan), or only in the irregular condition (‘Irreg’, magenta), or in both conditions (‘Both’, purple), or they swapped significant encoding of the respective joint from regular to irregular (‘J-Swap’, orange).

To examine encoding of forelimb kinematics on the population level we next used the random forest algorithm to predict forelimb joint movements based on the estimated spiking rates of all M1 L2/3 neurons within each imaging area (Fig. 5a,b; Online Methods). As a measure of predictive power we used the correlation of predicted and real joint motion. For both wheel conditions, the motion of all joints was significantly encoded by the local L2/3 populations for neuronal networks 1, 2, 3, 4, 6 and 7 (based on ROC-AUC ≥ 0.9 for true and shuffled distribution; see Online Methods and Supplementary Figs. 3 and 4). In neuronal network 5, prediction of elbow movements just missed significance and in neuronal network 9, only prediction of finger movements in the regular context and prediction of shoulder movements in the irregular context were significant. In neuronal network 8, none of the joint angles were predicted significantly even though shoulder in both conditions and elbow in the irregular condition came close (AUC = 0.88 in each case). When the predictive power in both conditions was compared, the same neuronal networks displayed significantly increased mean prediction for shoulder and wrist motion when the animals moved in the irregular instead of the regular context (Fig. 5c; *P* = 0.0075, t = −4.5492, *P* = 0.0845, t = −2.4140, *P* = 0.0288, t = −3.3834 for shoulder, elbow and wrist, respectively, paired t-test with *P* adjusted according to HB, df = 8, n = 9).

**Figure 5.**
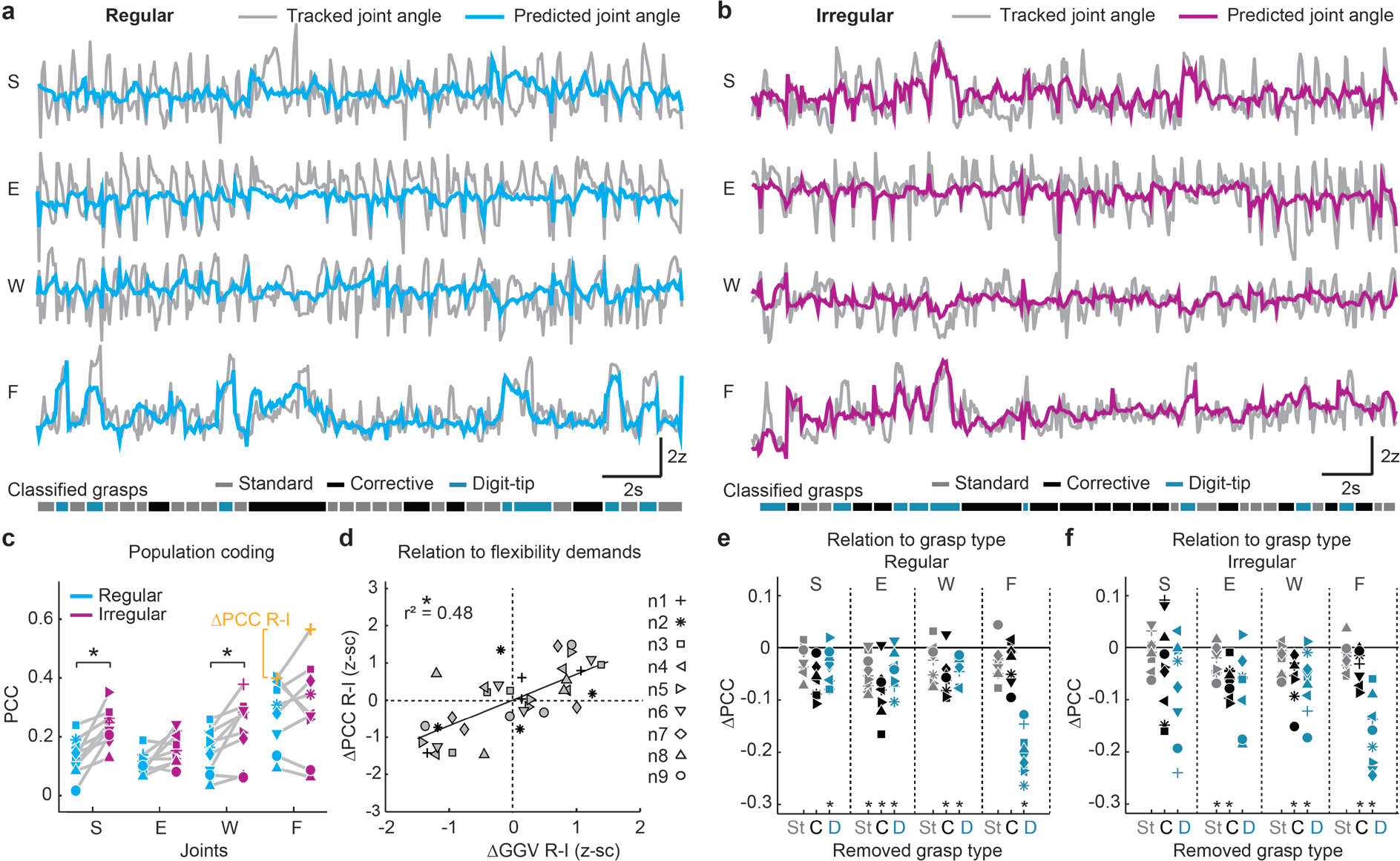
Prediction of forelimb variables from L2/3 activity. (**a**) Tracked shoulder (S), elbow (E), wrist (W) and finger base (F) joint angles (grey) for an example running period on the regular wheel together with angle changes predicted from the imaged L2/3 neuronal population activity using random forest regression (cyan). Bottom: Grasp classification based on kinematic criteria as shown in Fig. 1. (**b**) Example running period on the irregular wheel for the same neuronal population as shown in (a). Angle changes predicted from the imaged L2/3 neuronal population activity are depicted in magenta. (**c**) Forelimb joint movement prediction from activity of neuronal populations in M1 L2/3 in the regular (cyan) and irregular context (magenta), based on Pearson’s correlation coefficient (PCC) between real and predicted joint angle traces. Results are shown for all 9 recorded neuronal networks. Black asterisks indicate significantly different population coding between the regular and irregular condition for the respective joint; *P < 0.05 after adjustment according to Holm-Bonferroni, paired t-test. (**d**) Between-condition differences in prediction (ΔPCC) versus between-condition differences in grasp-to-grasp-variability (ΔGGV). ΔGGV (see also Fig. 1) was computed to quantify differences of joint flexibility demands in-between the regular and irregular context. Linear regression with clustered standard error (robust). (**e**) Population coding of each joint angle when one of the three grasps is removed from the dataset in the regular condition (St: Standard grasps, C: Corrective grasps, D: Digit-tip grasps). For each neuronal network, prediction changes (ΔPCC) are shown relative to the population coding when all grasps are included (zero line). (**f**) Same conventions as in (e), but for the irregular condition.

The predictive power for finger-base joint motion tended to surpass that of all other joints in the regular and irregular context but did not differ significantly between conditions (Fig. 5c, **Supplementary Video 3**; *P* = 0.7227, t = −0.3676). For all joints and both conditions, saturating population coding was in most cases achieved after inclusion of 20-40% of the population size (Supplementary Fig. 5).

Because one salient difference between both conditions is captured by the different flexibility demands of joint angle amplitudes (see grasp-to-grasp variability, GGV, in Fig. 1h), we next analyzed to what extent differences in predictive power in the regular versus the irregular context can be explained by context-dependent differences in GGV. We found that between-condition differences in joint angle prediction by neuronal networks were significantly explained by corresponding between-condition differences in joint angle GGV (Fig. 5d; *P* = 0001, r^2^ = 0.48, n = 9 networks, four joint angles for each; linear regression with clustered standard error; cluster variable = neuronal network which are thereby regarded separately; see Online Methods). Importantly, this dependence was true for all joints and not only for wrist and shoulder. Encoding for wrist and shoulder consistently increased in all neuronal networks from regular to irregular, as did the GGV, leading to respective significant differences. However, in neuronal networks 4, 5 and 9, for example, the encoding of finger movements decreased from regular to irregular, as did the GGV. As a consequence encoding and GGV of finger movements were not significantly different between conditions, but were positively correlated also in these networks.

Because of the large fraction of regular- and irregular-specific single-cell predictors of joint angles (see above) we next investigated to what extent the population coding of individual joint movements relies on context-specific neuronal configurations. When the random forest algorithm was trained in the irregular condition and joint movements then predicted for the regular condition, the predictive power significantly decreased for elbow and finger-base movements (Supplementary Fig. 6a; *P* = 0.0863, t = 1.9549, *P* = 0.0263, t = 3.4437, *P* = 0.0514, t = 2.7337, *P* = 9.2175*10^−6^, t = 11.8868 for shoulder, elbow, wrist and finger, respectively; paired t-test with *P*-values adjusted according to HB, df = 8, n = 9). Vice versa, when the random forest algorithm/decoder was trained in the regular condition and joint movements was predicted for the irregular condition, the predictive power significantly decreased for all joints (Supplementary Fig. 6b, *P* = 0.0168, t = 3.7541, *P* = 0.0111, t = 4.2582, *P* = 0.0156, t = 3.5225, *P* = 0.0125, t = 3.2072 for shoulder, elbow, wrist and finger, respectively; paired t-test with P-values adjusted according to HB, df = 8, n = 9). Together with the single cell analysis, these results imply that neuronal circuits in M1 L2/3 are reconfigured when the animal moves in different contexts. Overall, these findings suggest that neuronal networks of M1 L2/3 show enhanced motion encoding of those joint angles that demand higher grasp-to-grasp flexibility in a given contextual setting and vice versa.

We next asked how the encoding of individual joint movements by neuronal networks in M1 L2/3 relates to the three different grasp types. We therefore analyzed how the prediction of joint movements changes when one of the three grasp types was removed from the dataset (see Online Methods). This analysis showed that digit-tip grasps mainly contribute to the pronounced prediction of finger-base joints in both conditions, which decreased dramatically after removal of digit-tip grasps (Fig. 5e,f; *P* = 3*10^−6^, t = 13.7314, *P* = 0.0011, t = 6.7519 for regular and irregular condition, respectively). This result is in line with the high average activity of some cells during digit-tip grasps and their frequent additional encoding of finger movements (see above). Both in the regular and irregular context, all three grasp types contributed to a similar extent to the prediction of shoulder, elbow and wrist movements, even though their encoding tended to depend more on digit-tip grasps in the irregular condition (Fig. 5e,f).

### Context-dependent M1 encoding of ‘twin grasps’ with matching kinematics

Between conditions, we found two significant differences regarding the population coding of joint movements in M1 L2/3: First, the encoding of shoulder and wrist movements significantly increased in the irregular compared to the regular context. Second, context-dependent encoding differences could be explained by the corresponding contextual differences in joint flexibility demands. We therefore next asked to what extent these ‘between-condition effects’ might have been generated by differences in limb kinematics across conditions despite the occurrence of the same grasp types. Kinematic differences across conditions could possibly arise from the different fraction of grasps of each type in the regular and irregular context as well as from joint angle differences between grasps of the same type.

To probe a possible influence of kinematic differences between conditions, we investigated if the observed joint encoding differences and their explanation by differences in joint flexibility demands are preserved if only equivalent movements with matching joint angle kinematics on the regular and irregular pattern (‘twin grasps’) are considered. If preserved, at least part of the encoding differences must have emerged from the different contexts with regular or irregular rung spacing and cannot simply be explained by kinematic differences on the two types of wheels. We therefore compiled the most similar grasp pairs across conditions, separately for standard, corrective and digit-tip grasp clusters, based on the Euclidean distance of 4-dimensional joint angle vector pairs (Fig. 6a; see Online Methods). The selection procedure generated ‘twin’ standard, corrective and digit-tip grasp clusters that differed minimally across conditions (Fig. 6b,c). Re-analyzing only these twin grasps, we could confirm the differences found for joint motion encoding: Population encoding of shoulder and wrist motion was still significantly increased for the irregular compared to the regular wheel (Fig. 6d, *P* = 0.0275, t = −2.6910, *P* = 0.0399, t = −2.8978, for shoulder and wrist, respectively; paired t-test with *P* adjusted according to HB, df = 8, n = 9). In addition, between-condition differences in the encoding of all joints were still significantly explained by between-condition differences in their GGV (Fig. 6e; *P* < 0.0001, r^2^ = 0.5041, four joint angles for each of 9 neuronal networks; linear regression with clustered standard error; cluster variable = neuronal network). In fact, the variance of encoding differences explained by differences in GGV slightly improved compared to the whole dataset (r^2^ = 0.50 vs. r^2^ = 0.48), even though only the most similar grasping pairs and equal numbers of each grasp type were regarded. Moreover, we did not find any relationship between joint-angle dissimilarities within twin grasp pools and the observed encoding differences (Supplementary Fig. 7). Altogether, the results of the twin grasp analysis indicate that the observed joint angle encoding differences between the regular and irregular stepping pattern reflect distinct contextual needs for joint flexibility rather than merely kinematic joint movement differences between conditions.

**Figure 6.**
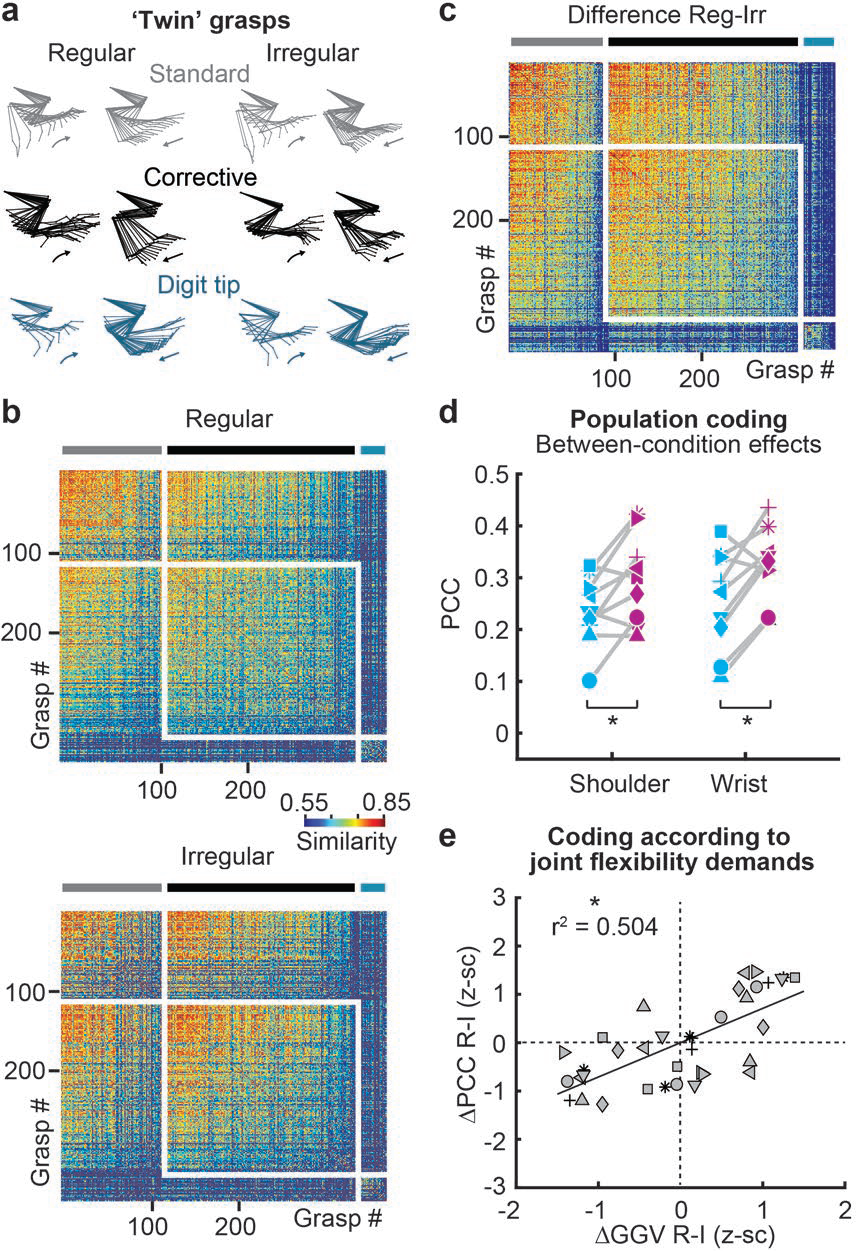
Encoding of forelimb variables and grasp types for twin grasps in different contexts. (**a**) Representative examples of twin grasp pairs for the regular and irregular condition. (**b**) Similarity matrices for the regular and irregular condition after the twin-movement pruning for the same example animal as in Fig. 1f: For each movement on the regular wheel, a near-identical “twin”-movement on the irregular wheel exists that belongs to the same grasp type and features minimal differences in the sample-point-wise Euclidean distance calculated for the 4-dimensional joint angle space. (**c**) Similarity matrix for the difference between regular and irregular after the twin-pruning. (**d**) Between the regular and irregular condition, the significant encoding differences for shoulder and wrist are preserved when only twin grasps are regarded. (**e**) Differences in the encoding of individual joint angles when grasps from each twin grasp cluster are pooled per condition versus differences in their total grasp-to-grasp variability from the regular to the irregular condition.

### Context-dependent encoding emerges during learning, increases precision of limb movements, and requires input from the higher motor area M2

We next asked how the observed M1 encoding of motion in individual joints according to their flexibility demands in the environmental context is related to motor learning, precision of limb movements, and interaction with the secondary motor cortex (M2). To probe the impact of M2 on contextual encoding in M1 L2/3 we conducted a subset of experiments, in which we injected a Cre-dependent hM4D(Gi)-construct into M2 and retrograde AAV-6 Cre virus into M1 of 2 additional mice, along with YC-Nano140 for calcium imaging (Fig. 7a, see also Online Methods). We were thus able to silence neurons projecting from M2 to M1 by injection of the otherwise pharmacologically inert synthetic ligand clozapine-N-oxide (CNO) ^52–54^. We recorded the activity of the neuronal networks 6 and 7 (mouse 6) as well as 8 and 9 (mouse 7) across the naive, learning and expert training phase as well as during the expert training phase while silencing M2-M1-projections (248 cells across 4 imaging areas, 62 ± 17.49 neurons per area). From the naive to the learning to the expert training phase, context-dependent encoding of flexibility demands consistently increased across mice and reached significance during the expert phase (Fig. 7d; r^2^ = 0.0036; *P* = 0.8051, r^2^ = 0.2446; *P* = 0.0719, r^2^ = 0.3648, *P* = 0.0348 for naive, learning and expert training phase, respectively; 4 joint angles for each of the 4 neuronal networks; linear regression with clustered standard error, cluster variable = neuronal network). During the expert phase, silencing of M2-M1-projections by injection of CNO considerably decreased context-dependent encoding when compared to vehicle injections (Fig. 7d). In fact, the amount of context-dependent encoding of flexibility demands subsided approximately to the level of the naive phase (r^2^ < 0.0001, P = 0.9891; 4 joint angles for each of the 4 neuronal networks; linear regression with clustered standard error, cluster variable = neuronal network).

**Figure 7.**
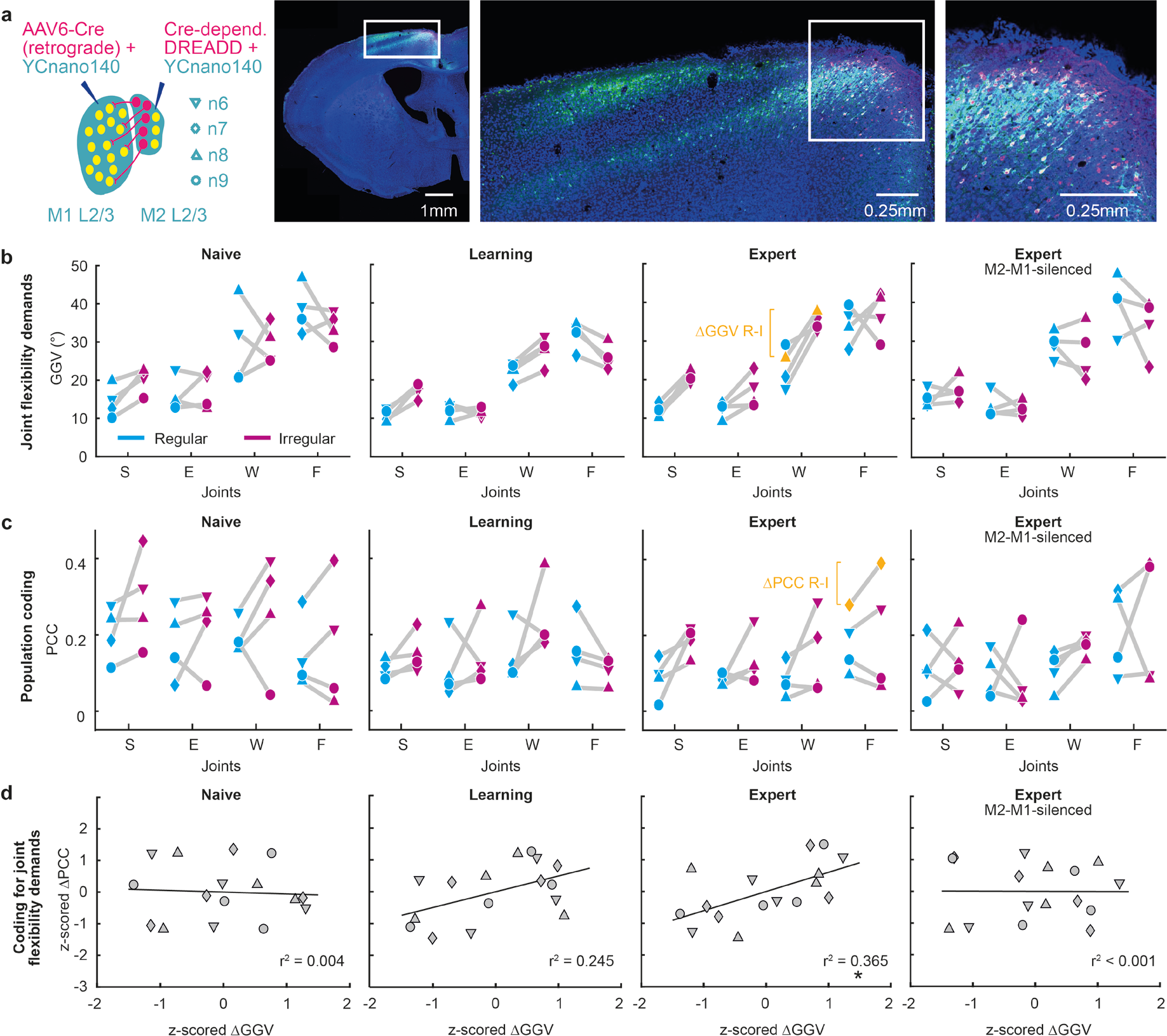
Emergence of context-dependent population coding of flexibility demands during learning and dependence on input from M2. (**a**) Expression of the DREADD system in M2 L2/3 and L5 (mCherry). Joint flexibility demands (**b**), population coding (**c**), and population coding according to joint flexibility demands (**d**) during ‘Naive’, ‘Learning’, ‘Expert’ and ‘Expert M2-M1-silenced’ phase, respectively. Note that context-dependent encoding according to joint flexibility demands emerges during learning and is disrupted after silencing of M2-M1 projections.

A single-cell coding analysis of all 248 recorded cells revealed that the fraction of neurons that predicted a particular joint in the regular or irregular context correlated inversely with the level of context-dependent encoding, decreasing from the naive (37%) over the learning (15%) to the expert training phase (11%) while increasing again during silencing of M2-M1-projections (40%; Fig. 8a). These results indicate that M2 increasingly suppresses redundant neuronal joint movement representation when the interaction with a new environment is learned.

**Figure 8.**
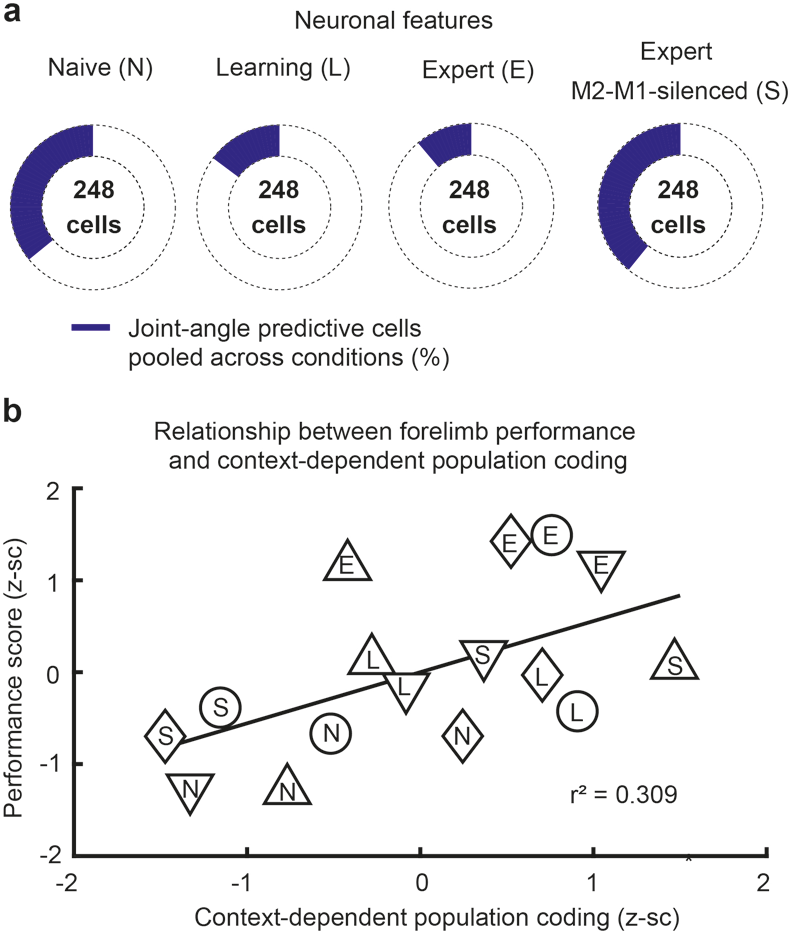
Neuronal features during training and M2-M1-silencing as well as context-dependent population coding versus grasp precision. (**a**) Total number of neurons that significantly predicted a joint during either condition (purple, as in Fig. 4) decreases from naive over learning to expert level and increases again in the expert M2-M1-silenced phase. (**b**) Context-dependent encoding of individual joint movements correlates positively with the precision of grasping actions, quantified by the forelimb performance score as shown in Supplementary Figure 1.

Finally, in line with the increased context-dependence of encoding during learning and its decrease upon silencing of M2-M1-projections, the forelimb performance score increased from naive to learning and to expert phase but substantially decreased following the disruption of M2-M1-input. Hence, the forelimb performance score for each neuronal network significantly correlated with the level of context-dependent encoding (Fig. 8b; r^2^ = 0.3091, P = 0.0467; 4 flexibility encoding indices for each of the 4 recorded neuronal networks; one for naive, learning, expert, and M2-M1-silenced phase, respectively; linear regression with clustered standard error, cluster variable = neuronal network). We conclude that context-dependent encoding of joint movements emerges during learning, entailing more accurate limb movements in the respective context, and that it requires intact information streams from the higher motor area M2.

## DISCUSSION

Our study extends the current concept of movement processing in the motor cortex network. Our main findings are that during motor learning neuronal networks in M1 L2/3 progressively encode joint movements according to their flexibility demands in the given environmental context. The context-dependent modulation of encoding occurs for equivalent movements, is conveyed by the higher motor area M2, and is paralleled by more precise limb movements. Encoding of joints according to contextual flexibility demands is accompanied by a low number of predictive neurons, thereby requiring low computational resources, and features context-specific neuronal configurations.

So far, neuronal activity in M1 during skilled locomotion on ladder paradigms has been mainly investigated in the cat motor cortex. These studies showed that M1 L5 neurons modify their firing rates during locomotion on ladders with regular or vertically displaced rungs^55^, during forelimb movement phases when evading an obstacle^9,10,56^, during expected and unexpected gait perturbations on a regular ladder^57^, and when accuracy demands are varied by comparing locomotion on flat surface, a regular ladder and regular ladders with different crosspiece widths^58,59^. However, none of these studies addressed the question how the required flexibility of individual limb joints in the given context impacts their coding in neuronal motor cortex networks. While two studies^58,59^ demonstrated different firing of M1 L5 neurons by comparing different accuracy demands, their tasks included a flat surface and regular ladders thus featuring virtually equivalent joint flexibility demands. Another study^55^ compared paradigms with different joint flexibility demands but neuronal activity was not related to motor output parameters and no context-variable such as required flexibility of joint movements was quantified.

Further studies probing the relationship between movement variables such as limb joints and neuronal activity in the motor cortex were performed in monkeys. In restricted, simplified movement sets, such as a set of reach directions, neuronal activity has been proposed to relate to direction, force, speed, joint angle, movement trajectories and muscle activity, among other variables^17,20–22,25,26,60,61^. In contrast, the representation of these variables was minor during naturalistic movements and much of the neuronal variance remained unexplained, even though neurons showed partial tuning to individual joint angles of preferred arm-postures^23,62^. One possibility may be that the difference of context-characterizing variables in natural behavior and restricted tasks contributed to the observed encoding differences since the contexts of natural behaviour and restricted tasks are likely to differ in demands such as the required flexibility of limb joints and presumably many others. So far the activity of neuronal networks in M1 has been predominantly related to kinematics or force of ongoing movements, and context-characterizing “meta”-variables may not have been acknowledged sufficiently. Context-dependent differential encoding of equivalent limb movements, as found here, suggests that incorporating contextual features into models of M1 encoding may contribute to a more coherent conception of motor cortex function.

Due to its anatomical position and connectivity, M1 L2/3 is a suitable candidate to integrate features of the current environmental context with motor commands. M1 L2/3 receives input from other cortical areas such as the somatosensory cortex^32,33^ and the secondary motor area M2^34–36^, which is thought to organize flexible motor behavior and to link relevant context information to motor processing, similar to the primate supplementary complex^37–39^. Neuronal populations in M1 L2/3 also send excitatory projections to M1 output neurons in layer 5^40,41^ whereby they can couple sensory information to motor output^42,44,63^. Our finding that silencing the M2 to M1 projections abolishes context-dependent modulation of movement encoding in M1 L2/3 suggests that M1 L2/3 learns to exploit contextual information from M2 to selectively route sensory information from those joints into corticospinal circuits, which require flexible re-adjustments in the given environment. Different routing of sensory information into the descending motor command during equivalent movements is likely to contribute to degeneracy in corticospinal circuits, which has been suggested to promote the generation of equivalent movements in many different contexts^64^. Purposeful coordination of specific and contextual sensory information to generate accurate movements in varying environmental conditions is also postulated by models of sensorimotor control^31^, and may refine the flexible recruitment of muscle synergies by corticospinal neurons that leads to appropriate limb movements during reaching or gait modifications^18,27,56,65^.

The concept that M1 tailors the representation of movement variables to contextual demands may also have important implications in clinical neuroscience. Considering context-dependent representations may for instance help to improve the performance of brain machine interfaces for the control of limb prosthetics in people with paralysis which remains challenging particularly with regard to environmental context changes^66^. Furthermore, this concept may contribute to the understanding of movement deficits in motor disorders as rodents with neurological M1 dysfunction due to a stroke or Parkinson lesion are known to lack the joint movement flexibility that is required during skilled locomotion especially across irregular rungs^3,45^. The requirement of intact motor cortex function for skilled locomotion on regular as well as irregular rungs is indicated not only by forelimb deficits after M1 lesions^45^ but also by our finding of impaired forelimb performance after silencing of M2-M1-projections. In this regard, our experimental paradigm also provides a novel approach that should be helpful for investigating cortical pathophysiology in rodent models of neurologic diseases with M1 dysfunction. A caveat of our paradigm is that we cannot exclude the possibility that the encoding of joint movements actually reflects a relationship with a motor variable that could be correlated with joint movements, such as for example muscle activity. M1 is known to represent both muscle activity and joint movement^22^, and our paradigm could therefore be advanced by supplementing the recording joint movement with implanted EMG-electrodes for mice^27^ in future studies.

In summary, we suggest that representation of limb movements in M1 significantly depends on the contextual environmental setting and not only on features that characterize the ongoing motor action. Learning which joint movements need to be varied with high flexibility in a given context and reinforcing their sparse representation might be a specific function of M1 L2/3 to focus control on the most relevant degrees of freedom in varying environments. Context-dependent encoding of limb movements in M1 L2/3 presumably emerges under the impact of higher motor areas during motor learning, entails precise limb movements and may therefore reflect a fundamental cortical processing strategy for adaptive motor behavior.

## METHODS

### Animal surgery and viral constructs

All experimental procedures were carried out according to the guidelines of the Veterinary Office of Switzerland and approved by the Cantonal Veterinary Office in Zurich. In 7 young adult (5-6 weeks) male transgenic ChR2 mice (Thy1-COP4/EYFP)^67^, we injected AAV2/1-*EF1α-YC-Nano140* (300 nl, approximately 1 × 10^9^ vg μl^−1^) into L2/3 of M1 (0.1 mm anterior, 1.9 mm lateral from Bregma, 300 μm below pial surface). Mice 1-5 were part of the first experimental series and mice 6-7 of the second experimental series. Mouse 6 and 7 were additionally injected with AAV-6 Cre-dependent hM4D(Gi)-mCherry virus into lower layer 2/3 as well as upper layer 5 of M2 (1.5 mm anterior, 0.5 mm lateral to Bregma, 500 μm below pial surface) and with AAV-6 Cre virus into M1 L2/3 (0.1 mm anterior, 1.9 mm lateral from Bregma, 300 μm below pial surface). hM4D(Gi) is a DREADD (“designer receptor exclusively activated by designer drug”)^68^ that we used in the second experimental series to chemogenetically silence M2 neurons with axonal projections to M1 L2/3, found mainly in lower L2/3 and upper L5^35^. hM4D(Gi)-DREADDs are activated by the otherwise pharmacologically inert synthetic ligand clozapine-N-oxide (CNO), resulting in membrane hyperpolarization and silencing of the infected neurons^52–54^.

It should be noted that recent studies pointed out potential caveats regarding the application of DREADD systems. While one group reported no evidence that CNO crosses the blood brain barrier^69^, this result is in contrast to the findings from another group^70^. One suggested possibility that activation of DREADDs in vivo is likely to be mediated by metabolism of CNO to clozapine^69^, which readily crosses the blood brain barrier, but has also affinity for serotonergic and dopaminergic receptors^71^. However, the affinity of clozapine for muscarinic-based DREADDs is substantially higher than for native receptors^68^. That the observed decline of forelimb performance in the expert M2-off phase was actually generated by effects of low metabolized clozapine doses on native receptors, is therefore in our opinion highly unlikely, especially since clozapine is known as anti-psychotic drug with minimal motor side effects^71^. Still, that the observed disruption of contextual flexibility encoding and the impairment of forelimb performance in the expert M2-off-phase was generated specifically by M2-M1-projections, must be regarded with the above mentioned reservations.

24 hours after virus injections, a circular cranial window (4-mm diameter in animals 1-5, 5-mm diameter in animals 6 and 7) was implanted over M1 around the injection coordinates^72^. Contralateral to the cranial window, an aluminium head post for head fixation (weight < 1 g) was implanted on the skull using dental cement. During the surgeries, mice were anaesthetized with isoflurane (4% induction, 2% maintenance). After the surgery the animals were treated for analgesia with Rimadyl (Carprofen; 5 mg/kg body weight, s.c.) as well as the antibiotic Rocephin (40 mg/kg body weight, s.c.) and returned to their home cage for recovery. For the following 3 days, Rimadyl and Rocephin were injected once per day and the animal’s well-being was evaluated at least twice per day for the first two days after surgery and at least once per day during the following 5 days.

### Behavioral setup and training

All mice were handled, habituated to head fixation, and at first trained to locomote on top of a 23-cm diameter wheel with a flat surface. Before each trial, a brake blocked the wheel and prevented the animals to initiate locomotion. After an auditory start cue (12-kHz and 16-kHz tones for regular and irregular pattern, respectively, or vice versa) and release of the brake, animals had to initiate skilled locomotion and cover a predefined distance of 15-30 cm (in one animal only 10-15 cm per run) until an auditory stop cue (8-kHz tone) indicated the end of the trial (‘run’). In successful trials, 2 seconds after the stop tone, the brake was reactivated and the animal received a reward of sweet water (2 µl). In unsuccessful trials, in which the mouse did not traverse the predefined distance within the given time period, the animal was punished with a time-out (~20 s) before the next run. After one week of training on the wheel with the flat surface animals were placed for the first time on the 23-cm diameter regular or irregular ladder wheel which were custom-built from two acrylic glass rings as well as carbon rungs and which emulated the rung ladder test for rodents^3,45,46,73^. For the regular wheel rungs were spaced at constant 1-cm distances and for the irregular wheel rung distances varied unpredictably between 0.5 to 3 cm. During locomotion on both wheels, mice continuously localized rungs before reaching actions through whisking. The task was designed so that the next rung is in reach of the whiskers. Due to the training week on the flat surface, all animals successfully ran the predefined distance according to the tone cues in more than 80% of the trials on the regular and irregular wheel. On each day, we then evaluated the first 10 successful runs on the regular and irregular wheel with the forelimb performance score^3,45^ which was then averaged across the 10 runs of each condition. Based on the forelimb performance score, the first 4 days were regarded as ‘naive’ training phase, days 5-8 were regarded as ‘learning’ training phase and days 9-12 were regarded as ‘expert’ training phase in which the forelimb performance score reached a plateau (Supplementary Fig. 1). In the first subset of experiments (animals 1-5), calcium imaging was performed in one M1 L2/3 neuronal network per mouse (networks 1-5) during the expert phase. In the second subset of experiments (animals 6 and 7), calcium imaging was performed in two M1 L2/3 neuronal networks per mouse (networks 6-9) during the ‘naive’, ‘learning’ and ‘expert’ phase as well as additionally during the expert phase after M1-projecting M2-neurons were silenced by injection of CNO (‘expert M2-M1-silenced’ phase, corresponding to days 13-16, see Supplementary Fig. 1). To allow the comparison between the expert M2-M1-silenced phase and the three training phases, mice 6 and 7 received vehicle injections during the expert, learning and naive training phases. For all 9 neuronal networks, the sequence of the regular and irregular condition was randomized.

### Limb motion tracking

To allow the analysis of forelimb kinematics during light stimulation and calcium imaging of M1, the skin overlying defined anatomical landmarks of the right forelimb was shaved and tattooed with a commercially available tattooing kit (Hugo Sachs Elektronik, Harvard Apparatus GmbH). On the forelimb, we marked the vertebral border of the scapula along with shoulder, wrist, metacarpophalangeal joint (MCP; also referred as ‘finger-base’ joint throughout the manuscript), and the tip of the third digit. Limb kinematics during the optogenetic light stimulation was tracked at 30-Hz frame rate with a video camera (Logitech B910 HD) monitoring the right side of the animal. To allow tracking of limb kinematics during calcium imaging, the right side of the animals was illuminated with two 940-nm infrared LED light sources and recorded at 90-Hz frame rate (1280 × 640 pixels) using a high-speed CMOS camera (A504k; Basler). For analysis, time series of kinematic variables were downsampled to the imaging frame rate (18 Hz) using cubic spline interpolation as implemented in MATLAB.

The markers on the skin of the forelimb were semi-automatically tracked offline, frame-by-frame using the ClickJoint 6.0 software (ALEA solutions GmbH), extracting two-dimensional coordinates (*x* for horizontal, *y* for vertical) for every marker and time point^46^. Based on these coordinates, the software modeled limb segments as rigid straight lines between markers and calculated the angles in each joint for consecutive frames. To minimize artifacts caused by skin stretching over the elbow joint, the position of the elbow was deduced from the shoulder and wrist coordinates as well as from the upper (~1.1 cm) and lower (~1.2 cm) forelimb length^46^. For subsequent analyses, we considered the angle changes in the shoulder, elbow, wrist and MCP joints. Hand movement was quantified by the *x*- and *y*-coordinates of the MCP joint in each video frame and the reaching distance was calculated from *x* and *y* by Pythagorean addition.

### Optogenetic motor mapping

Two weeks after window implantation mice were anesthetized with ketamine-xylazine (100 mg kg^−1^ ketamine, 10 mg kg^−1^ xylazine) for optogenetic motor mapping^47^. Mice were placed in a hammock with all four limbs dangling freely (Fig. 2a). Using a stereoscope with a motorized scanning system, a 473-nm laser beam was directed to 100 spots in mice 1-5 and 225 spots in mice 6 and 7 on the left motor cortex. The spots were arranged in a 10×10 grid in mice 1-5 and in a 15×15 grid in mice 6 and 7 (square area of 9.66±0.55 mm^2^, n = 7 mice). Each of the 100 or 225 spots was hit in random order and stimulated for 500 ms at 100 Hz (pulse duration 4 ms; laser intensity < 100 mW mm^−2^). In all animals, the beam diameter was adjusted to 130 µm at the level of the motor cortex in the window center by using a reference micrometer grid. During stimulation of M1, the right side of the animal was monitored with a camera for subsequent offline-analysis of forelimb kinematics. The angle changes in the shoulder, elbow, wrist and finger-base joint during light stimulation of M1 were quantified using ClickJoint 6.0 software (ALEA Solutions GmbH). Spatial maps of joint angle changes were spline interpolated to 145×145 pixels in mice 1-5 and 225×225 pixels in mice 6 and 7. Subsequently, the half-maximal (50%) contours were calculated in MATLAB. The stimulation spot, which caused the maximal combined absolute angle changes in shoulder, elbow, wrist and finger joint, was selected as “forelimb focus region” for calcium imaging.

### Two-photon calcium imaging

In the first experimental series, calcium imaging was performed with a custom-built two-photon microscope controlled by HelioScan^74^, equipped with a Ti:sapphire laser system (100-fs laser pulses; Mai Tai HP; Newport Spectra Physics), a water-immersion objective (16×CFI75 LWD, NA 0.80, Nikon), galvanometric scan mirrors (Cambridge Technology), and a Pockel’s cell (Conoptics) for laser intensity modulation. In the second experimental series, a custom-built two-photon resonant-scanning microscope controlled by ‘Scope’ (http://rkscope.sourceforge.net/)^51^ was used in conjunction with a Mai Tai HP DeepSee laser (Spectra-Physics). For calcium imaging, YC-Nano140 was excited at 820 nm to avoid simultaneous activation of ChR2 in dendrites of L5 neurons (see below). Fluorescence was collected in epi-collection mode with 480/60 nm (CFP) and 542/50 nm (YFP) emission filters using photomultiplier tubes. Image series were acquired at 18 Hz with 128×64 pixel resolution (galvanometric system) and at 21.768Hz with 942×362 pixel resolution (resonance system).

Calcium imaging data from YFP and CFP channels were imported into MATLAB for subsequent processing steps. Lateral motion in both data channels were corrected with the TurboReg algorithm^75^. Individual neurons were selected manually from the mean image of each single-trial time series as regions of interest (ROIs). The background-subtracted mean pixel value of each ROI was extracted for both channels and applied to express neuronal calcium signals as relative YFP/CFP ratio change ΔR/R = (R − R_0_)/R_0_ in which we employed a sliding window across the dataset to infer the baseline ratio R_0_. To yield an estimate of instantaneous spiking rate (SR), calcium signals were deconvolved using a Wiener filter algorithm assuming an exponential kernel as single-action potential evoked ΔR/R transient (amplitude 4.54%, decay time constant 0.673 s, onset time constant 0.186 s)^49^. The smoothness parameter was set to 0.01.

To identify single neurons that displayed activity related to particular grasp types (Fig. 4b) we first calculated the mean SR traces across all grasps for each type (traces normalized in duration). A neuron was considered significantly responsive for a particular grasp type if the SR value of the mean trace, averaged over the entire grasp duration, surpassed the mean + 4 s.d. of the distribution of average SR values obtained from shuffled neuronal SR traces (corresponding to a chance level of 0.003%; 500 times shuffling of the grasp order). SR traces were also used for subsequent correlation and population coding analyses (see below).

### Electrophysiology of L5 neurons

The use of transgenic ChR2 mice is advantageous for performing calcium imaging in motor cortex areas identified using optogenetic mapping, which does not require mechanical tissue perturbation with electrodes. To ensure that subsequent two-photon imaging in L2/3 did not affect the activity of ChR2-expressing L5 neurons through depolarization of their apical dendrites, we performed cell-attached recordings of L5 neurons in 8 additional transgenic ChR2 mice (Fig. 2d and e). Blind juxtacellular voltage recordings were obtained from putative L5 neurons using glass pipettes (4–7 MΩ resistance) filled with control extracellular solution (in mM: 145 NaCl, 5.4 KCl, 10 HEPES, 1 MgCl_2_, and 1.8 CaCl_2_) and an Axoclamp 2B amplifier (Molecular Devices), preamplified, and digitized at 20 kHz with an ITC-18 board (InstruTECH) controlled by custom-written IGOR Pro software (WaveMetrics). Positive pressure (20-30 mbar) was applied while navigating the pipette in the tissue with a micromanipulator (Luigs & Neumann) to approach neurons. ChR2-expressing L5 neurons were identified by the pronounced spiking rate increases induced by blue (488 nm) laser light stimulation through an optical fiber placed a few millimeter above motor cortex (fiber output power about 11 mW). The effect of two-photon excitation was assessed by imaging in L2/3 above the recorded L5 neuron using the same experimental settings as in our calcium imaging experiments (near-infrared [NIR] light of 820-nm wavelength; illumination power < 45 mW). Following two-photon imaging in L2/3, blue-light stimulation of M1 was repeated to confirm that the neuron was still spiking. We analysed the number of spikes evoked for the four conditions (“Blue light on”, “Blue light off”, “NIR light on”, “NIR light off”) with spikes detected with a threshold routine: Spikes were assigned to those time points when the voltage difference crossed a threshold of 7 s.d. above mean baseline. Statistical significance was tested by paired t-tests between each of the 4 conditions with post-hoc adjustment of p-values according to Holm-Bonferroni (HB). Paired t-tests were applied after the Anderson-Darling test was used on the paired differences between each of the 4 conditions to test for normality.

### Analysis of grasping actions

Classification of grasping actions into the three grasp types was performed using custom-written functions in MATLAB (Version 7, MathWorks). We first defined single grasp cycles based on local minima found in the horizontal *x*-component of the reaching distance vector. To distinguish full grasp cycles from corrective grasping actions that typically occurred in front of the reach (creating subcycles with secondary maxima), only minima with values lower than the mean of all local minima and maxima were accepted. The number of subcycles during each grasp cycle was then defined as the number of local maxima of the Savitzky-Golay-filtered reaching distance vector (taking the larger number for *x*- and *y*-component). In addition, the mean finger extension was computed for each grasp cycle. Based on these measures we classified each grasp cycle into one of three types according to the following criteria:

- standard grasp: one subcycle, mean finger extension < 170°
- corrective grasp: two or more subcycles, mean finger extension < 170°
- digit-tip grasp: one or several subcycles, mean finger extension > 170°

Grasp amplitude *A* for each joint was calculated as the difference between the maximum and minimum of angle position. The grasp-to-grasp variability (GGV) for a particular joint angle (JA) and condition *C* (regular or irregular wheel) was defined as the mean absolute amplitude difference between each grasp and its preceding grasp per run, averaged across all runs:

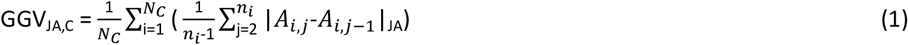

Here, *A*_i,j_ denotes the amplitude of the *j*-th grasp of the *i*-th run, *n*_i_ the number of grasping cycles during the *i*-th run, and *N*_C_ the number of runs for condition *C*. A high GGV value indicates that the movement amplitude of this particular joint was frequently substantially changed from one grasp to the next. In contrast, low GGV values indicate rare and little grasp-to-grasp adjustments of motion amplitude. The GGV of a grasp sequence thus reflects the requirement for grasp-to-grasp adjustments set by the specific movement context (here regular or irregular rung pattern).

To quantify the similarity of pairs of grasping actions we first normalized (z-scored) all kinematic JA variables (S-shoulder, E-elbow, W–wrist, F-finger base). We then resampled kinematic traces for all individual grasps via interpolation to a fixed number of sample points (*N*_fix_ = 160) in order to align all grasps with a normalized duration. As distance measure we calculated the sum of sample-point-wise Euclidean distance *d* for pairs of 4-dimensional resampled JA vectors **p** and **q**:

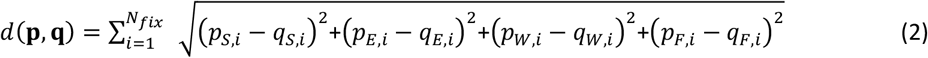

where *i* runs over all sample points. A problem with this definition is that two discrete grasping vectors featuring nearly the same time course of coordinated JA angle changes, but slightly temporally shifted would yield an artificially high Euclidean distance. To reduce this problem, we used dynamic time warping^76^ to allow temporal warps, coupled for all joint angles, over a restricted time window. Maximally allowed time warps were 33% of the grasp duration, i.e. 53 sample points. Optimal time warping for each **p** and **q** vector pair was found by minimizing *d*(**p**,**q**). Finally, to bound similarity values between 1 (maximum similarity) and 0 (maximum dissimilarity) the similarity value *S* was calculated as

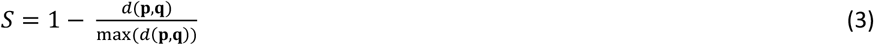

Grasp-similarity matrices were clustered according to the grasp-type classification and within each cluster sub-sorted according to similarity values with respect to the mean grasp for the respective grasp type.

### Motion prediction using the random forest algorithm

We used the random forest algorithm (RFA)^77^ to predict limb motion, i.e., kinematics of shoulder, elbow, wrist and finger-base joints, and grasp types from the activity of either single neurons or neuronal populations. The RFA is a multivariate, non-parametric machine learning algorithm and utilizes bootstrap aggregation of regression trees. We adopted the Treebagger function implemented in MATLAB and specified 150 trees and default settings for minimum leaf size and number of variables to select at random for each decision split. These parameters were an appropriate trade-off between computation time and decoding accuracy. For the prediction of joint angle kinematics a regression RFA was used.

After concatenation of normalized (z-scored) joint angle traces of all grasps for one condition (regular or irregular wheel), the algorithm was trained to predict the real joint angle changes from the instantaneous SR traces of one (single neuron prediction) or all neurons of the recorded network (population coding) on a randomly selected subset of grasps, comprising 70% of the dataset (training set). For cross-validation, the trained algorithm was then evaluated on the remaining 30% of the dataset left out during training (test set). To quantify the predictive power, we computed the Pearson’s correlation coefficient (PCC) between joint angle changes predicted by RFA in the test set and the corresponding real joint angles. We repeated this procedure 500 times, thereby obtaining a distribution of 500 ‘true’ predictions, that came from 500 randomly selected test sets of one ‘true’ dataset. As shuffled control, the ‘true’ assignment of calcium traces to motor output parameters in each trial was randomly shuffled between all trials for 500 times. For each shuffling, training (70%) and corresponding test set (30%) were randomly selected and the predictive power was quantified as for the ‘true’ dataset. This process generated a second distribution of 500 shuffled predictions that came from 1 randomly selected test set of 500 shuffled datasets. This procedure allowed us to compare the lowest predictive power from all test sets in the true dataset to the highest predictive power from different test sets in different shufflings (high predictive power in the shuffled distribution could arise due to the random data-shuffling itself and additionally due to the random selection of a certain test set with high predictive power; therefore, comparison of these shuffled and true distributions is far stricter than for example comparing the mean prediction of 500 shufflings with the mean prediction of the true dataset which is also an accepted approach). We then applied a ROC-analysis on the shuffled and true distribution and calculated the area under the ROC-curve (ROC-AUC). According to Hosmer and Lemeshow ^78^, ROC = 0.5 indicates no discrimination, 0.7 ≤ ROC-AUC < 0.8 indicates acceptable discrimination, 0.8 ≤ ROC-AUC < 0.9 indicates excellent discrimination and ROC-AUC ≥ 0.9 indicates outstanding discrimination. We regarded the predictive power of a single neuronal network for a particular joint angle as significant if the AUC was ≥ 0.9. Since the analysis was more frequently applied in the single cell analysis (493 neurons), the predictive power of a single cell for a particular joint angle was considered significant if the AUC was ≥ 0.95. The ROC-analysis does not require that the two distributions are normally distributed. Even though each neuron predicted a particular joint to the highest degree, a considerable amount of predictive power was sometimes also observed for further joints (Supplementary Fig. 8). If a single neuron displayed significant prediction for more than one joint, the identity of a neuron (e.g ‘finger-predictive cell’) was determined by the joint that was predicted to the highest degree.

As single, summarizing value for the predictive power in the true and shuffled distributions, we defined the mean PCC. To test the significance of differences in predictive power between the regular and irregular condition with regard to all animals, paired t-tests or Wilcoxon signed rank tests with post-hoc adjustment of *P*-values according to Holm-Bonferroni were used for the mean prediction values of the 9 neuronal networks (Regular vs. Irregular). The decision if a paired t-test or a Wilcoxon signed rank test was applied depended on the result of a previous Anderson-Darling test that tested normality of the paired differences between the 9 neuronal networks. In the regular and irregular condition, neuronal population RFA prediction was applied for the whole dataset (Fig. 5c) as well as after removal of standard, corrective and digit-tip grasps (Fig. 5e and f). In this analysis, we randomly selected an equal number of grasps (corresponding to the number of grasps in the smallest cluster in each network and condition) from the standard, corrective and digit-tip grasp cluster to avoid training bias of the random forest algorithm. We then computed the population coding as described above for the combined grasps from the three clusters as well as after removal of one grasp type. The change in predictive power (ΔPCC) was then calculated by subtracting the predictive power of the pool with all three grasp types from the predictive power of the respective pools with two grasp types. To compare the underlying encoding rules between the regular and irregular condition, we also trained the RFA on a given neuronal network in one condition (e.g. regular) and used this training set to predict the motor output parameters in the other condition (e.g. irregular).

To explore the relationship between the activity of single neurons and grasp types, the activity trace of each neuron was averaged across the respective grasp type, after time-normalization for each grasp. A neuron was regarded as grasp-type related if the mean of its averaged activity surpassed the mean + 4 s.d. of the distribution of averaged means after shuffling of the grasp order (500 shufflings, chance level 0.003%).

### Correlation between differences in encoding and in grasp-to-grasp variability

We investigated whether between-condition-differences in prediction of individual joint angles relate to between-condition-differences of their grasp-to-grasp variablitiy (GGV). We first calculated for each neuronal network and joint angle the differences in prediction (PCC) and GGV (irregular minus regular), yielding 4 ΔPCC and 4 ΔGGV-values per neuronal network (one for each joint). We then z-scored the 4 values for ΔPCC and ΔGGV for each animal. Then, we calculated a linear regression with clustered standard error (regression in Stata, type clustered robust, animals as cluster variable) of ΔPCC differences versus ΔGGV differences pooled for all mice (Fig. 5d and 6d). A significant positive linear relationship here indicates that between-condition increases in GGV of individual joint angles are accompanied by between-condition enhancements of their encoding in M1 L2/3 neuronal networks and vice versa. For example, GGV for the shoulder angle increased from the regular to the irregular condition, as did its encoding in neuronal networks of M1 L2/3. Importantly, linear regression with clustered standard error regards the regression separately for each neuronal network and has therefore stricter requirements for significance than a simple linear regression with values from all neuronal networks pooled.

### Twin grasp analysis

Similarity of grasp pairs across the two conditions (regular and irregular) was quantified as described above using Euclidean distance of normalized JA vectors in 4-dimensional JA space (Equ. 3). Starting with the standard grasp cluster, we first selected the most similar grasp pair across conditions. The respective standard grasps were then no longer available for further selections. From the remainder of grasps in the standard grasp clusters, we again selected the most similar pair and repeated this procedure until all grasps in the standard grasp cluster of one condition were consumed. The same procedure for ‘twin grasp’ selection was performed on the corrective and digit-tip grasp clusters. After this selection procedure, each twin cluster in the regular condition featured the same number of grasps as its counterpart in the irregular condition. The mean deviation of all joint angles in twin grasp pairs over the grasp duration was 13.17° ± 0.98° across animals (mean ± SD, see Supplementary Fig. 7 for values with regard to individual neuronal networks and individual joints). In the twin grasp analysis, the grasp-to-grasp variability of each joint angle was calculated as before for all grasps in the regular or irregular condition (not only for twin grasps) because we regarded this measure as a general feature of the entire movement sequence and its environmental context.

### Statistics

All statistical analyses were computed in MATLAB R2017a and Stata 14. Data are shown as individual data points for each observational unit. To compare two data sets (e.g. regular vs. irregular), paired t-tests (two-tailed) or the Wilcoxon signed rank test were used. The decision if a paired t-test or a Wilcoxon signed rank test was applied depended on the result of a previous Anderson-Darling test that tested normality of the paired differences between the 9 neuronal networks. Linear regression with clustered standard error was calculated in Stata 14 (linear regression, type clustered robust). Linear regression with clustered standard error is sensitive to the regression in each observational unit (in our case values of the respective neuronal network) and therefore stricter than a simple linear regression of values that have been pooled across neuronal networks.

For all statistical tests, the post-hoc Holm-Bonferroni method for multiple comparisons was applied by adjusting the *P*-value correspondingly, and the respective exact *P*-value is given in the *Results* section. A significant difference between two data sets was assumed when the Holm-Bonferroni-corrected *P*-value was below 0.05 (indicated by one asterisk in figures).

## ACKNOWLEDGEMENTS

We thank M. Wieckhorst, H. Kasper and S. Giger for technical assistance, C. von Achenbach and A. Brändli for help with data analysis, B. Seifert for statistical advice, A. Caflisch for discussions, and A. Banerjee, L. Egolf and A. Gilad for comments on the manuscript. This work was supported by the Swiss National Science Foundation (SNSF) (grants 310030-127091 and 31003A-149858 to F.H), the EU-FP7 program (PLASTICISE project 223524 to M.E.S. and F.H.), and the Dr. Wilhelm Hurka Foundation.

## AUTHOR CONTRIBUTIONS

W.O designed the study under supervision of F.H.. W.O. performed optogenetic mapping and calcium imaging experiments of the first and second experimental series, in the latter together with P.S.. W.O., P.S. and A.S.W. trained the animals. W.O., H.L., F.H. analyzed the data. W.O., P.S., A.S.W. and L.S. performed tracking of joint angles. L.S. did perfusions of the animals, histology and immunohistochemistry. I.C. performed the electrophysiological control experiments. M.v.H., P.B. and F.V. provided technical support. W.O. and F.H. wrote the paper.

---

**Supplementary Figure S1.**
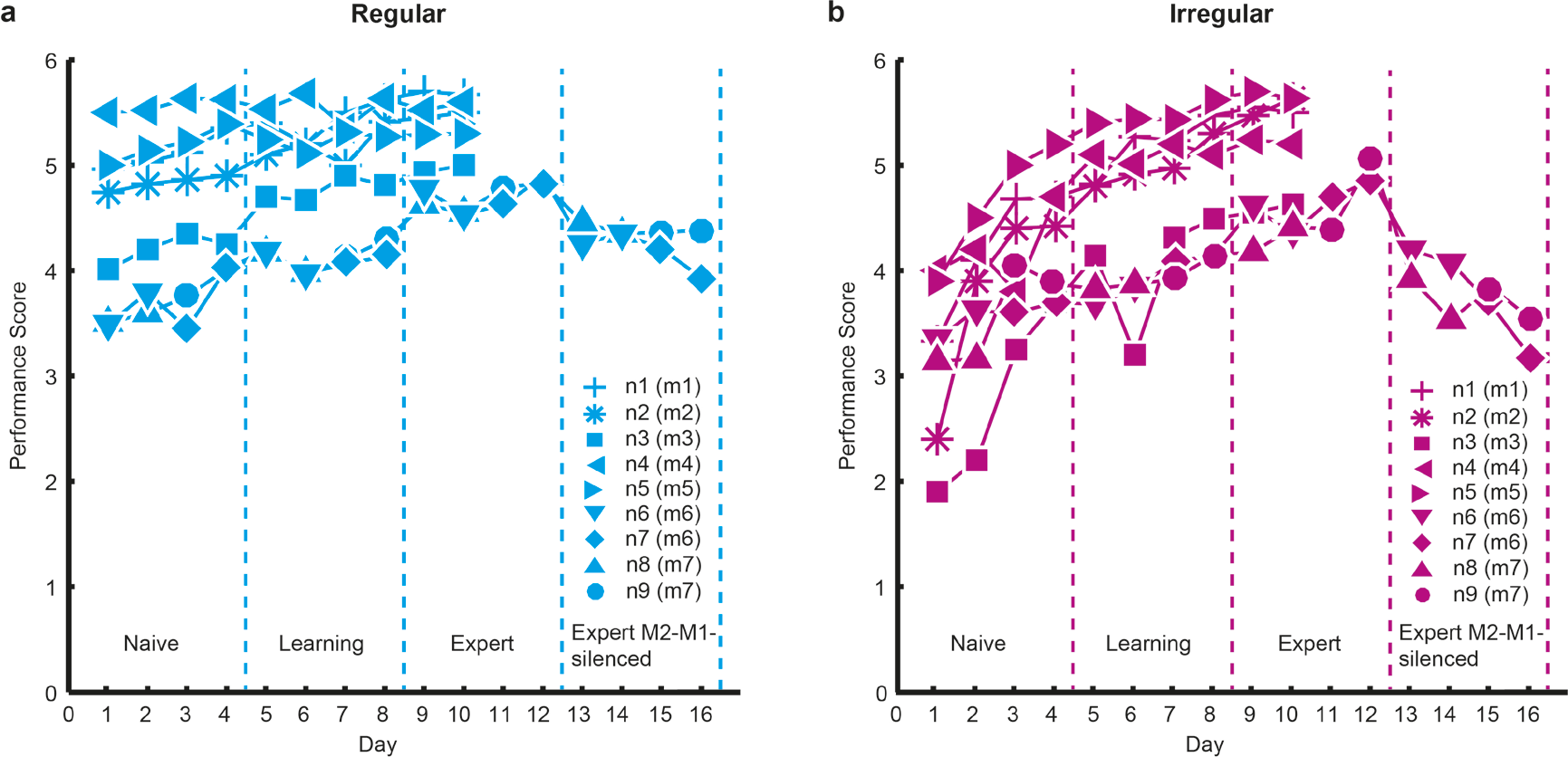
Forelimb performance score during learning of the motor task. (**a**) Evolution of the forelimb performance score for individual mice (n = 7) during learning of skilled locomotion and silencing of M2-M1-projections on the regular pattern (cyan). m1 – m7 refers to mouse 1 to 7, n1-n9 to neuronal network 1 to 9; (**b**) Evolution of the forelimb performance score for the same mice during learning of skilled locomotion and silencing of M2-M1-projections on the irregular pattern (magenta); based on the performance score we divided motor learning during both conditions into ‘naive’ (days 1-4), ‘learning’ (days 5-8) and ‘expert’ phase (days 9-12, performance score at saturating level); in mice 1 to 5 (1^st^ subset of experiments), calcium imaging was only applied during the expert phase; in mice 6 (n6 and n7) and 7 (n8 and n9), which correspond to the 2^nd^ subset of experiments, we included an additional expert phase with silencing of M2-M1 projections (‘Expert M2-M1-silenced’), and calcium imaging was performed during all phases. Rating according to the forelimb performance score: 0 = Total miss; 1 = Deep slip; 2 = Slight slip; 3 = Replacement; 4 = Correction; 5 = Partial placement; 6 = Correct placement.

**Supplementary Figure S2:**
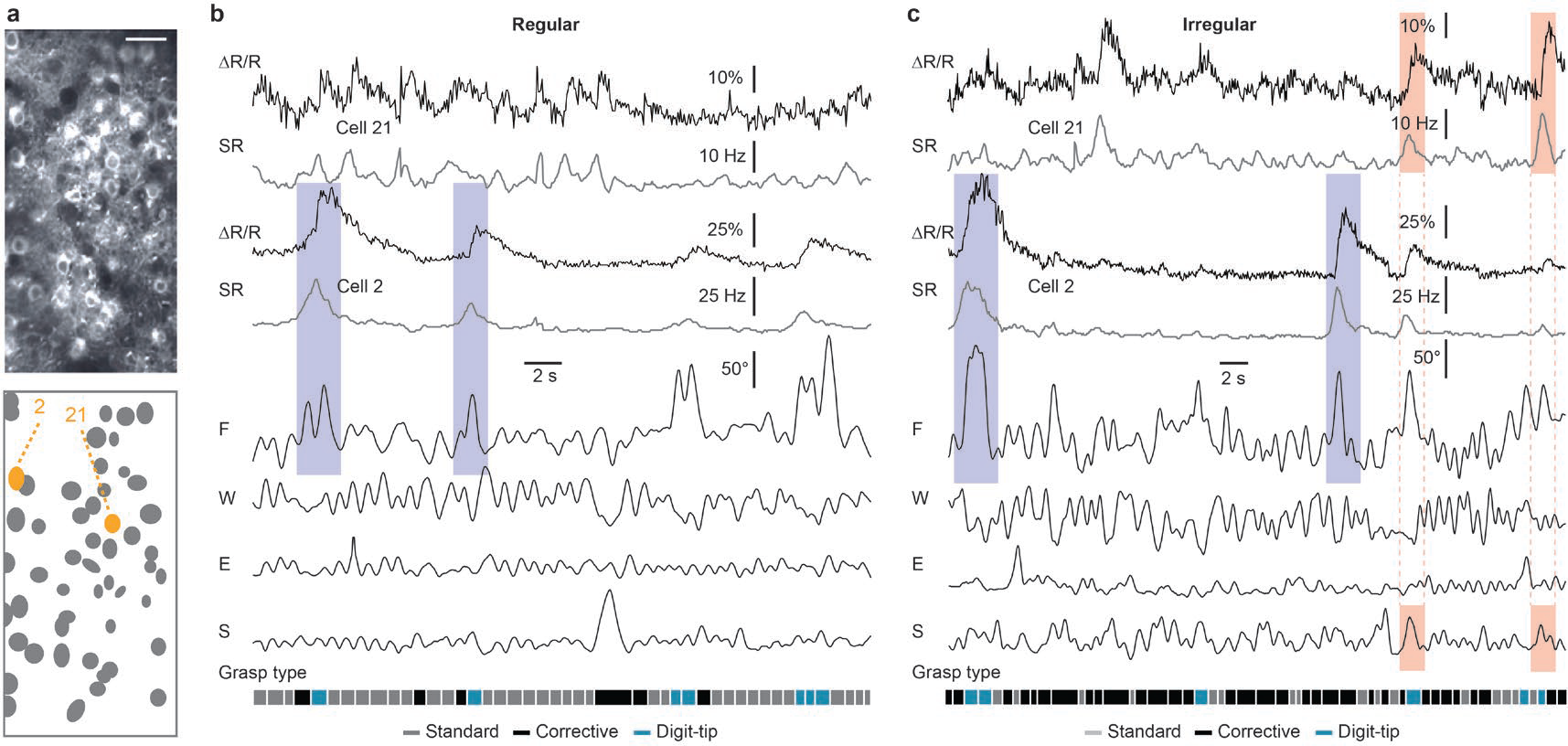
Calcium imaging data and movement variables in a different mouse compared to Figure 3. (**a**) Upper panel: Two-photon image of the YC-Nano140-expressing neuronal poplation that was measured with calcium imaging during skilled locomotion. Lower panel: Schematic of the selected regions of interest, with neurons 2 and 21 marked in orange. (**b**) Regular condition: Raw ΔR/R calcium signals (black traces) and deconvolved spiking rates (SR, dark grey traces) for the two cells marked in a, along with simultaneously recorded joint angles and the classified grasp types. (**c**) Irregular condition: Same conventions as in (b). Salient neuronal responses to finger base movements are indicated by the blue shaded areas. Salient neuronal responses to shoulder movements are highlighted by red shaded areas.

**Supplementary Figure S3:**
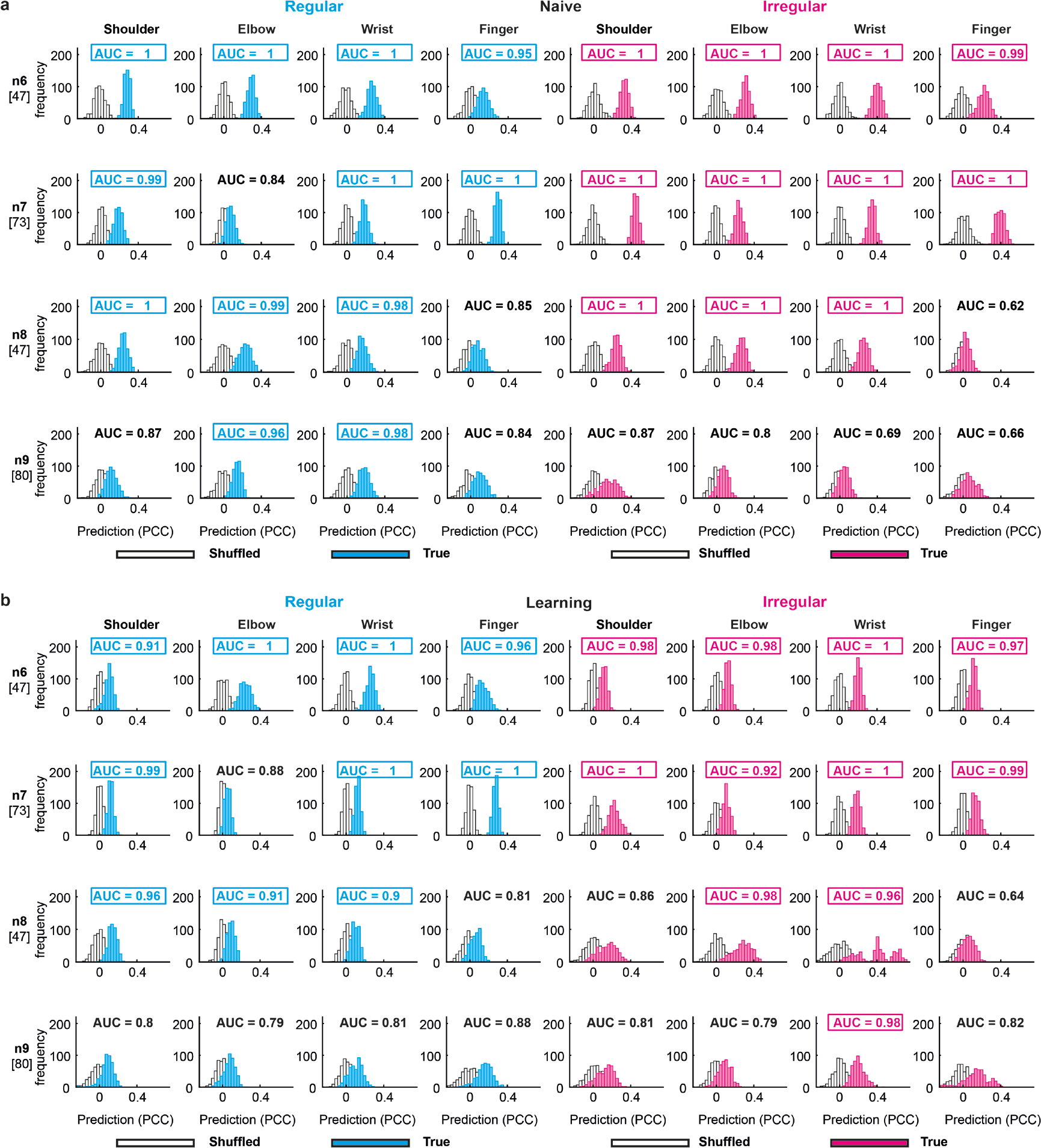
True and shuffled encoding performance during naive and moderate phase. Naive training phase: Prediction histograms: One 30% test set was randomly selected from 500 shuffled datasets (white for regular and irregular) or 500 30% test sets were randomly selected from the one true data set (grey for regular, black for irregular). The 2 distributions of 500 predictions in each recorded neuronal network (n6-n9, number in square brackets below corresponds to the cell count of each neuronal network) during the naive (a) and learning training phase **(b)** are compared by calculating the area under the curve (AUC) of a ROC-analysis. AUC-values ≥ 0.9 in the respective neuronal network are regarded as significant encoding. Note that high encoding in the shuffled distribution can arise as a consequence of the random shuffling and additionally by random selection of the test set. Still, true and shuffled distributions are in some cases completely separated.

**Supplementary Figure S4:**
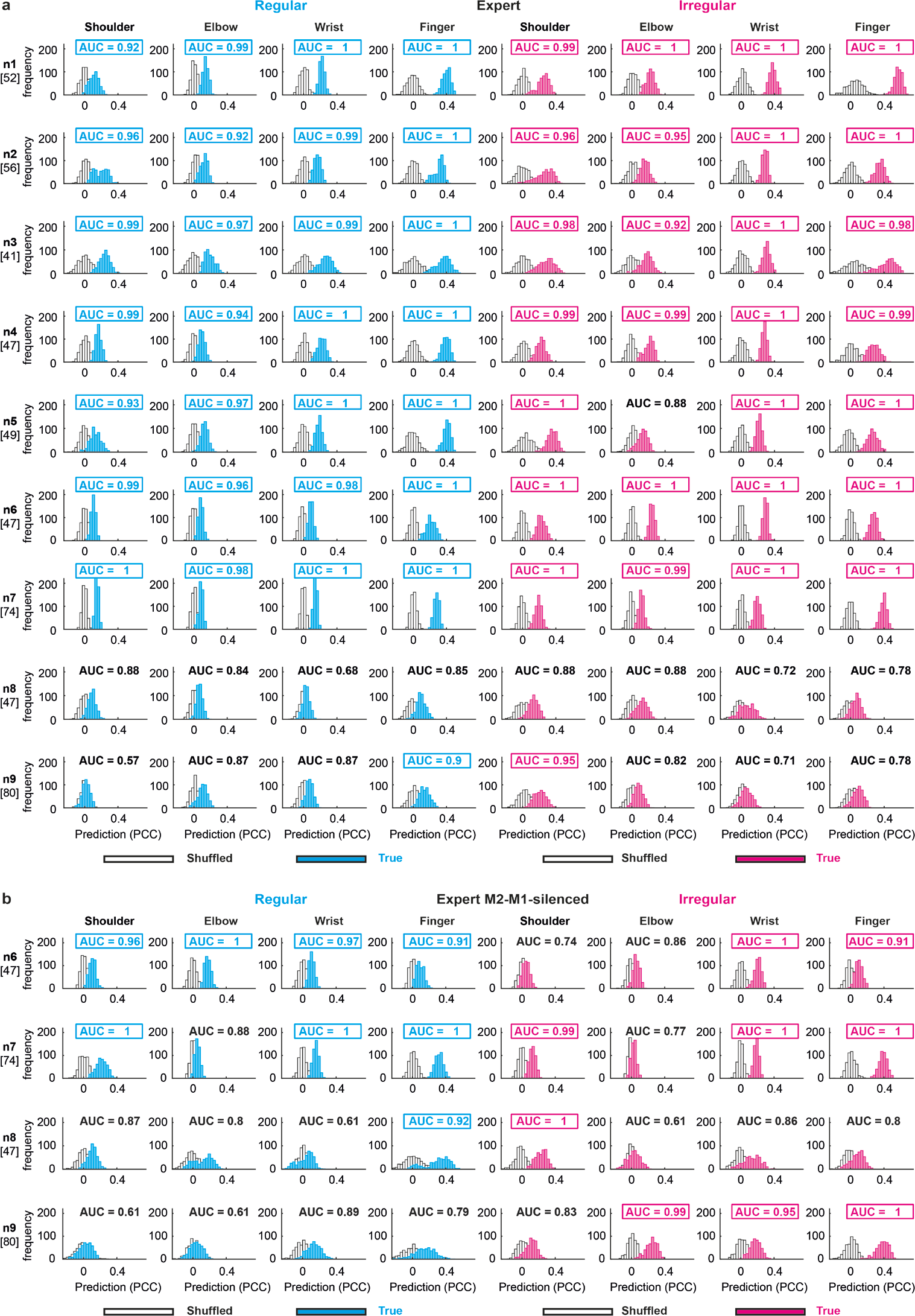
True and shuffled encoding performance during expert and expert M2-M1-silenced-phase. Same analysis procedure and conventions as in Supplementary Figure 3. **(a)** Encoding of forelimb joint movements in M1 L2/3 during the expert training phase. Altogether 9 neuronal networks (n1-n9, number in square brackets below corresponds to the cell count of each neuronal network) in M1 L2/3 have been recorded. **(b)** Encoding of forelimb joint movements in M1 L2/3 during chemogenetic silencing of M2-M1-projections using the DREADD-construct (“expert M2-M1 silenced phase”).

**Supplementary Figure S5:**
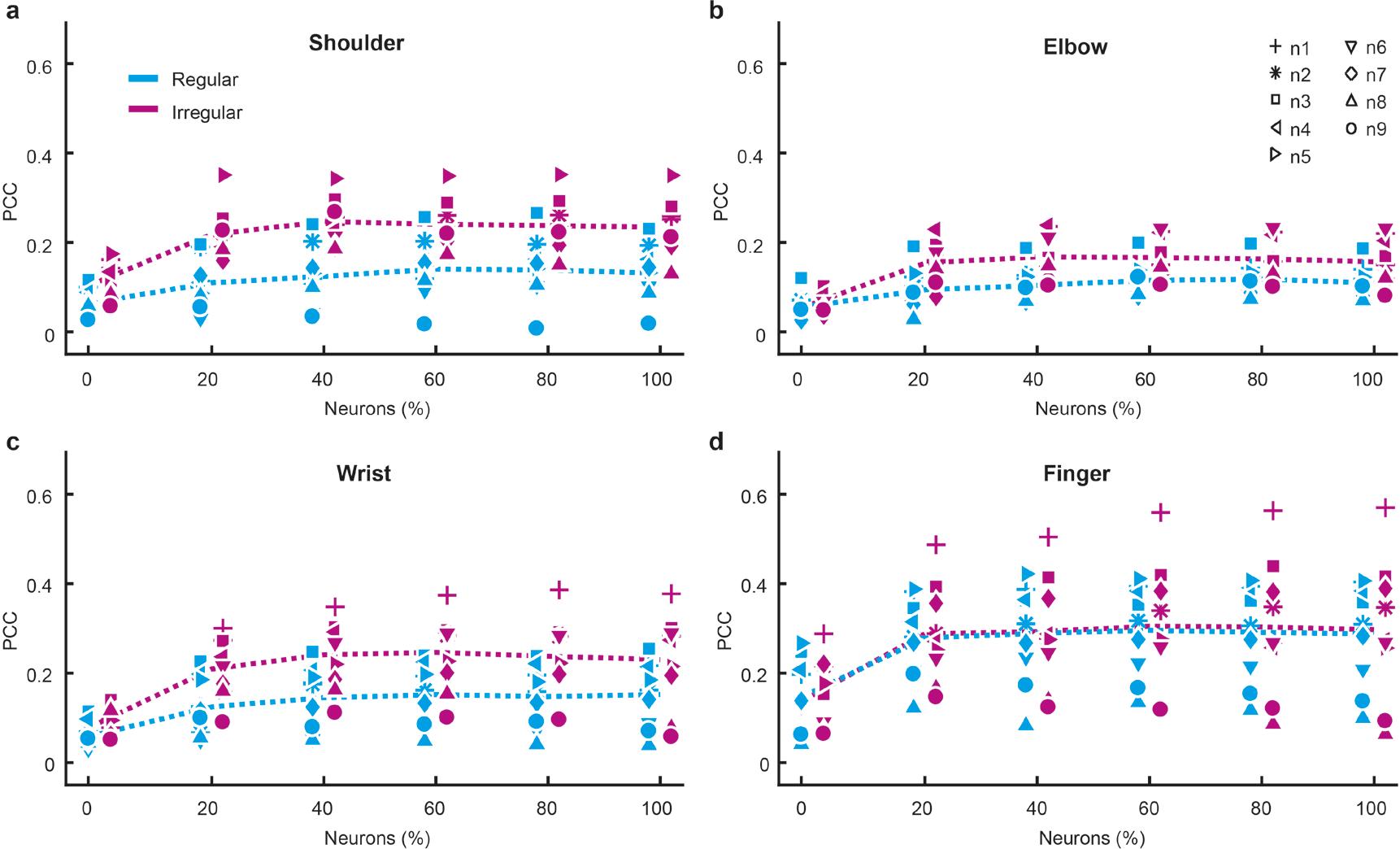
Prediction accuracy of forelimb joint angles as a function of population size. (**a**) Prediction of the shoulder joint angle as a function of population size, quantified by the Pearson correlation coefficient (PCC) between real and predicted joint angle traces and pooled across animals; for each neuronal network, cells with decreasing prediction strength have been added to the population coding, starting with best joint-angle-predictive single cell. Cyan symbols: Regular. Magenta symbols: Irregular. Dashed cyan curve: Mean for the regular condition. Dashed magenta curve: Mean for the irregular condition. (**b**), (**c**), and (**d**) Data for elbow, wrist and finger joints, respectively. Same conventions as in (a). Saturating prediction accuracy is mostly achieved after inclusion of 20-40% of the population size.

**Supplementary Figure S6:**
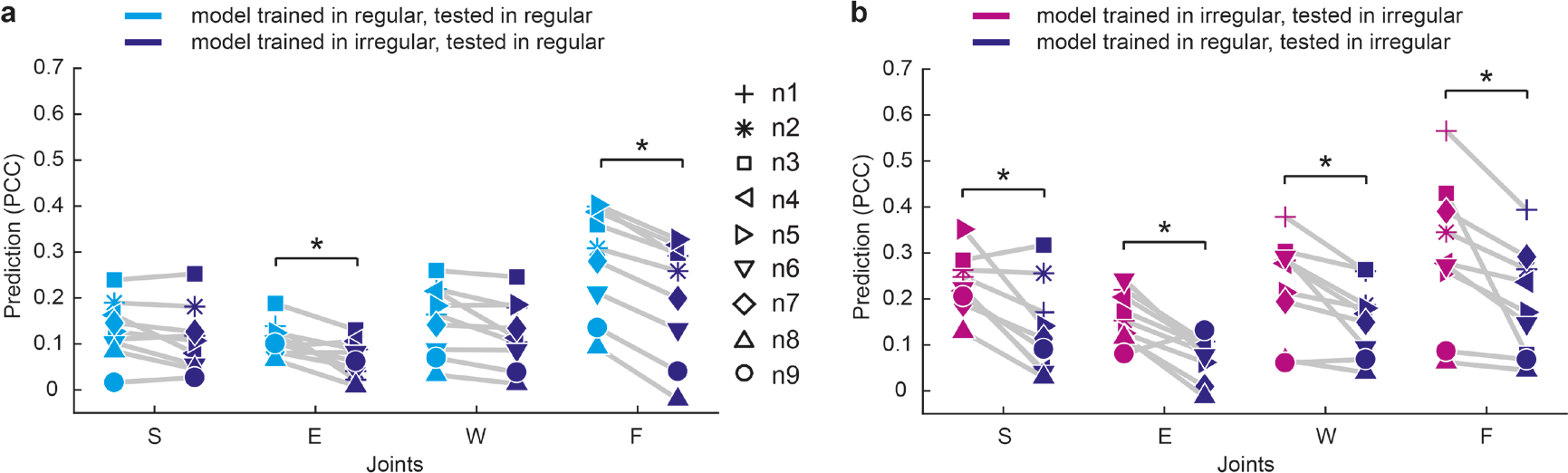
Decoding decrease when random forest is trained in the opposite condition. (**a**) Original decoding in the regular context (cyan) and decoding in the regular context if the prediction model was created from the same neuronal network in the irregular condition (purple; 70% training sets on the irregular condition, 30% test sets in the regular condition). Prediction power significantly decreases for elbow and finger. (b) Original decoding in the irregular context (magenta) and decoding in the irregular context if the prediction model was created from the same neuronal network in the regular condition (purple; 70% training sets in the regular condition, 30% test sets in the irregular condition). Prediction power significantly decreases for all joints.

**Supplementary Figure S7:**
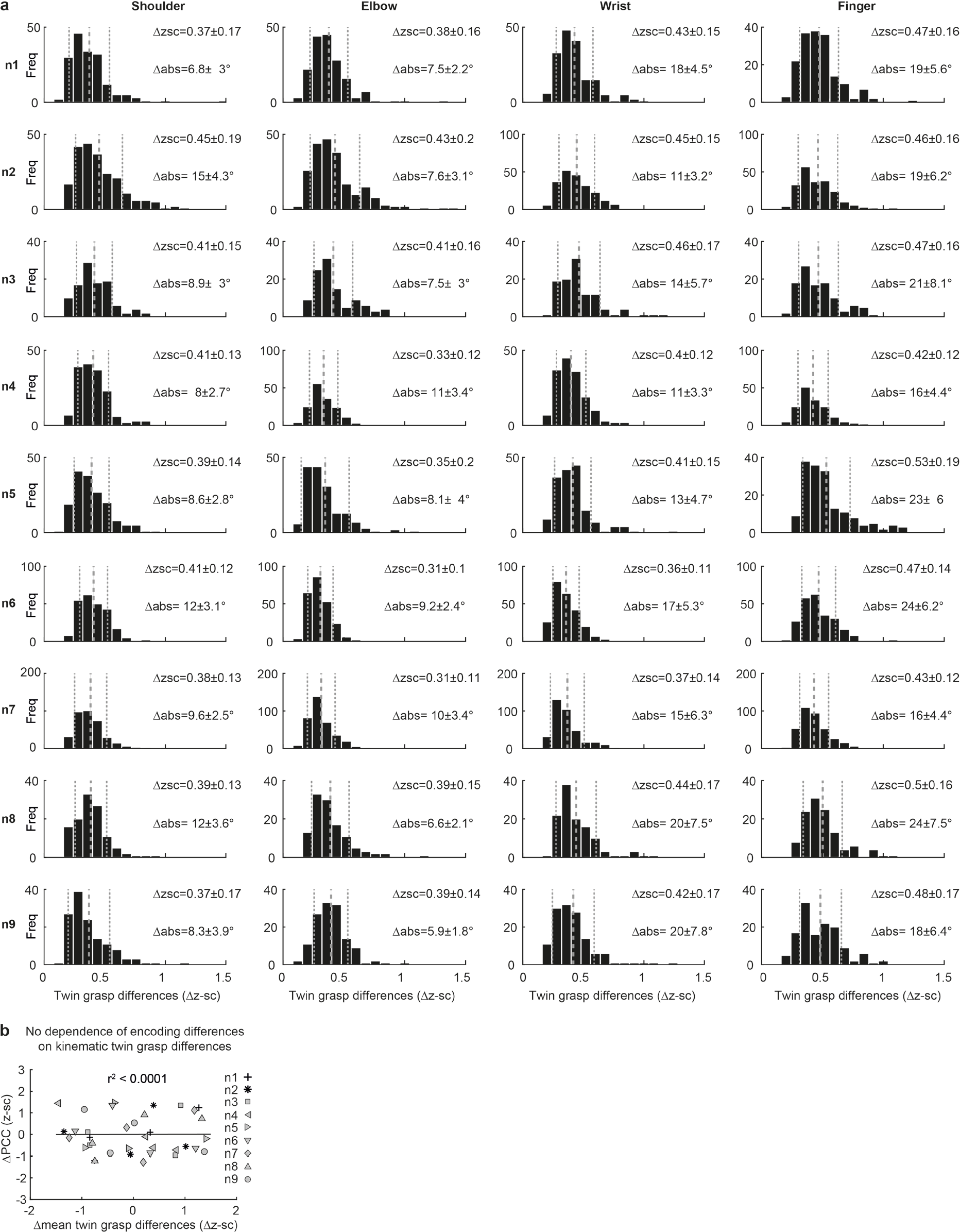
Joint angle (JA) differences between twin grasps and relationship with encoding differences. **(a)** Histograms of JA differences between all twin grasps for regular and irregular. Distributions show differences of z-scored JAs, and respective mean and SD are visible in each plot, along with mean and SD of twin grasp differences when JAs were not z-scored. (**b**) Relationship between encoding differences (ΔPCC) and mean JA differences (Δz-sc) for each joint during twin grasps. Note that encoding differences of JAs during twin grasps do not correlate with the dissimilarity of JAs during twin grasps. Linear regression with clustered standard error (robust, cluster variables = neuronal networks).

**Supplementary Figure S8:**
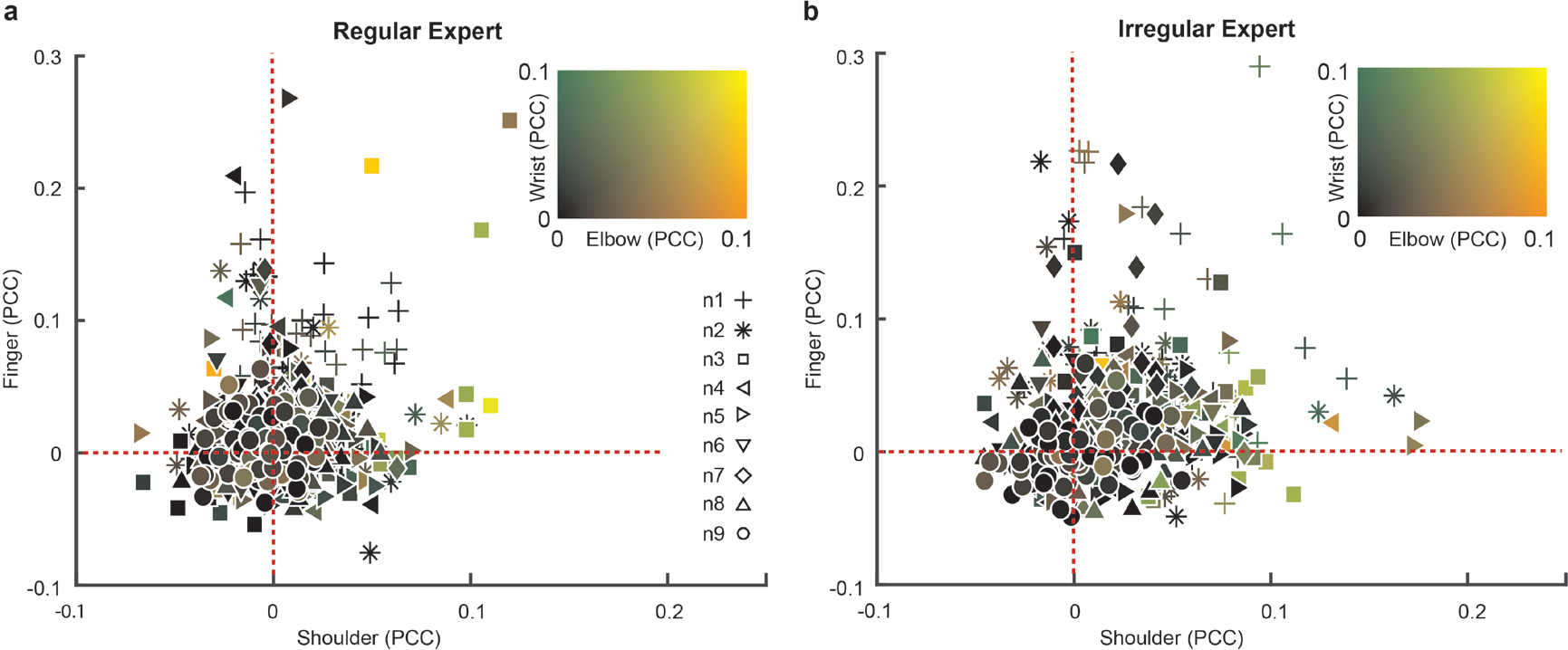
Prediction of forelimb joint angles using the activity of individual cells. (**a**) Four-dimensional plot showing the prediction of forelimb joint angles from the activity of individual cells, based on the Pearson correlation coefficient (PCC) between real and predicted joint angle traces in the test sets of the cross-validation procedure. Prediction values from all neurons of the 9 recorded neuronal networks are shown for the regular pattern. For each cell, the prediction for the shoulder and finger angles can be concluded from the spatial location in the diagram while the prediction for elbow and wrist can be derived from the color code in the upper right corner. (**b**) same conventions as in (a) but for the irregular condition. Each symbol represents data of a different neuronal network; n1 – n9 refers to neuronal networks 1 to 9. Note that negative correlation means that the relationship of this neuron to the respective joint was opposite between training and test set. Therefore, the cross-validated correlation values of cells that significantly encode a joint in the complete dataset, are always positive in this plot.

**Supplementary Video 1. Classification of three different grasp types.** Stick-figure movies of forelimb movements showing individual grasping actions (grey) along with the mean grasping action (green), separately for each of the three classified grasp types. To facilitate the illustration, each grasping action is normalized in time from start to end of the grasp. Data from one example mouse.

**Supplementary Video 2. Neuronal population activity during skilled locomotion on regular and irregular rung ladders.** Videos illustrating skilled locomotion (left panels) and simultaneously recorded time series of two-photon calcium imaging data for a L2/3 neuronal population in M1 (right panels). The upper row displays data acquired when the mouse was running on the regular wheel, the lower row when the same mouse was running on the irregular wheel. ⍰R/R values were measured in the same neuronal network under the two conditions and are overlaid in red pseudocolor code. Data from one example mouse. Scale bars 50 ⍰m. Frame rate 18 Hz. Replay 1.5x real time.

**Supplementary Video 3. Prediction of individual joint motion based on neuronal population activity.** The upper row displays data for the regular wheel, the lower panel for the irregular wheel. Left panels: Stick-figure movie of forelimb grasps showing the real movements of individual joints (grey) and the cross-validated prediction based on the combined activity of all neurons in the recorded network (purple) with the shoulder point affixed. Middle panels: Bar graph illustrating the temporal evolution of the Pearson’s correlation coefficients (PCC) between real and predicted joint angles accumulated over the time period shown. Right panels: Subplots showing the same grasping actions as in the left panels, but with each of the four joints affixed.

